# Metformin Stabilizes the Abdominal Aorta in Aneurysm by Restoring VSMC Mitochondrial Homeostasis via the AMPK–SIRT1–PGC-1α Axis

**DOI:** 10.64898/2026.02.27.708352

**Authors:** Baoyu Gao, Yuehuan Zeng, Ye Lu, Xinyu Yuan, Dongping Yang, Shaojun Lin, Jun Zhou, Buwen Liang, Yaliang Jie, Suping Ding, Jiahe Xie, Zhidong Yuan

**Author notes:** Corresponding authors (Z. Yuan). Corresponding authors (J. Xie). Corresponding authors (S. Ding).

## Abstract

**BACKGROUND:** Abdominal aortic aneurysm (AAA) is a life-threatening condition with >80% mortality upon rupture and no effective pharmacotherapy available. Despite epidemiological evidence linking metformin use to reduced AAA progression, its mechanism remains elusive. Notably, peroxisome proliferator-activated receptor γ coactivator 1α (PGC-1α, encoded by *Ppargc1a*) is downregulated in human AAA, yet its functional role in metformin’s protection is unknown.

**METHODS:** We employed porcine pancreatic elastase (PPE)-induced murine AAA, VSMC-specific Ppargc1a knockout (*Ppargc1a*^VSMC-KO^), primary VSMC senescence models, and pharmacological inhibition (Compound C for AMPK; Ex-527 for SIRT1) to define the AMPK–SIRT1–PGC-1α axis.

**RESULTS:** Metformin significantly inhibited AAA expansion, suppressed VSMC senescence (p53/p21↓, SA-β-gal↓), and preserved contractile phenotype (SMTN↑, IL-6/TNF-α↓). Crucially, all benefits were abrogated in *Ppargc1a*^VSMC-KO^ mice, which exhibited accelerated aneurysm growth, mitochondrial fragmentation, ATP depletion, and ROS accumulation. Mechanistically, metformin activated AMPK/SIRT1 to upregulate PGC-1α; AMPK or SIRT1 inhibition blocked this cascade and reversed protection.

**CONCLUSION:** Metformin restrains AAA by restoring VSMC mitochondrial homeostasis via the AMPK/SIRT1→PGC-1α axis, positioning PGC-1α as a non-redundant, cell-autonomous guardian against vascular degeneration. These findings provide a mechanistic foundation for repurposing metformin and developing PGC-1α-targeted therapies in AAA.

## 1. Introduction

Abdominal aortic aneurysm (AAA) is a life-threatening degenerative vascular disease characterized by localized, irreversible dilation of the infrarenal abdominal aorta, clinically defined as a maximum diameter ≥ 3.0 cm or ≥ 50% expansion relative to the adjacent normal segment^1^. AAA typically progresses asymptomatically, yet its ruptured carries a mortality rate of 80–90%. Globally, AAA rupture is estimated to cause 150,000–200,000 deaths annually^2,3^. Epidemiological data indicate a prevalence of 4-7% in men over 65 years and 1-2% in age-matched women^2,4^. Major risk factors include smoking, hypertension, hyperlipidemia, and atherosclerosis^5^. Although open surgery and endovascular aneurysm repair (EVAR)^6^ are effective for high-risk patients, no pharmacological therapy is currently approved to halt AAA progression—underscoring an urgent unmet clinical need for targeted intervention strategies.

Intriguingly, clinical studies consistently report a lower incidence and slower expansion of AAA in patients with type 2 diabetes mellitus (T2DM), with further risk reduction observed among those receiving long-term metformin therapy^7,8^. As a first-line antihyperglycemic agent^9^, metformin exerts pleiotropic effects beyond glycemic control—including anti-aging, anti-inflammatory, and mitochondrial protective actions^10^. Multiple cohort studies and preclinical models have confirmed an inverse correlation between metformin use and AAA development^11–13^, suggesting that its protective effects may operate via glycemia-inependent mechanisms.

Cellular senescence and pathological phenotypic switching of vascular smooth muscle cells (VSMCs) are pivotal drivers of AAA pathogenesis^14–16^. As the predominant cellular component of the aortic media, VSMCs exhibit remarkable plasticity. Under chronic stressors such as oxidative damage and inflammatory cytokines, they shift from a contractile phenotype (characterized by high expression of smoothelin [SMTN] and SM22α) to a synthetic/secretory phenotype (inflammatory microenvironment: elevated expression of IL-6, TNF-α) ^17–19^. This transition compromises vascular wall integrity not only by weaking contractile function but also by promoting extracellular matrix degradation and paracrine inflammation—ultimately culminating in medial degeneration and aneurysmal dilation.

Peroxisome proliferator-activated receptor γ coactivator 1α (PGC-1α) serves as a master regulator of mitochondrial biogenesis and energy metabolism. By coactivating transcription factors such as NRF1/2 and ERRα, PGC-1α orchestrates the expression of genes involved in oxidative phosphorylation, antioxidant enzymes (e.g., SOD2), and fatty acid oxidation—thereby sustaining mitochondrial fitness and redox homeostasis^20^. Mitochondrial dysfunction—manifested as ATP depletion and excessive reactive oxygen species (ROS) accumulation—is a well-established upstream trigger VSMC senescence^21,22^, positioning PGC-1α as a critical modulator of this process. Notably, PGC-1α expression is significantly downregulated in both human and mouse AAA tissues^23,24^, implicating its potential role in VSMC homeostasis disruption. Nevertheless, direct evidence linking PGC-1α to the regulation of VSMC senescence, phenotypic switching, and metformin-mediated protection remains lacking.

To address these gaps, we employed a porcine pancreatic elastase (PPE)-induced murine AAA model, complemented an angiotensin II (Ang II)-stimulated primary VSMC senescence system, to systematically investigate: (1) whether metformin attenuates AAA progression by suppressing VSMC senescence; (2) the functional necessity and mechanism contribution of PGC-1α in this process; and (3) whether the AMPK/SIRT1 signaling axis mediates metformin-induced of PGC-1α upregulation. Our findings delineate a novel “metformin → AMPK/SIRT1 → PGC-1α → mitochondrial homeostasis → VSMC stability” regulatory axis, offering both mechanistic insights and promising therapeutic targets for pharmacological intervention in AAA.

## 2. Materials and Methods

### 2.1 Experimental Animals

All experiments were performed using male C57BL/6J mice (10–12 weeks old, 25–30 g) and genetically modified strains including *Ppargc1α*^flox/flox^, *Rosa*26^mTmG^, and *Itga8*-*CreER*^T2^ mice, which were housed in the SPF-level barrier facility of Gannan Medical University under a 12-hour light/dark cycle with ad libitum access to food and water. Animal care and experimental procedures were approved by the Biomedical Research Ethics Committee of Gannan Medical University (Approval No.: 2022201) and conducted in accordance with relevant guidelines. Wild-type C57BL/6J (Jiangsu Huachuang Xinnuo Medical Technology Co., Ltd.), *Ppargc1α*^flox/flox^ and *Rosa*26^mTmG^ mice were obtained from commercial vendors (Cyagen Biosciences Inc.), while *Itga8*-*CreER*^T2^ mice were kindly provided by Dr. Lv Qing (Chongqing People’s Hospital, China). Mice were randomly assigned to experimental groups.

### 2.2 Construction of Murine AAA Model

Following the methodology described in reference^25^, a mouse AAA model was established by incubating porcine pancreatic elastase (PPE) externally on the renal abdominal aorta wall. Briefly, mice were weighed and administered 0.5% sodium pentobarbital (0.1 mL/10 g, intraperitoneal injection). Thirty minutes prior to surgery, butorphanol (0.1 mg/kg) was administered subcutaneously (SC). The abdomenal region was shaved and disinfected. A midline abdominal incision was made extending from the xiphoid process to the bladder to expose the infrarenal abdominal aorta. A small piece of gelatin sponge was placed on the exposed aortic segment. After aspirating excess moisture, 30 µL of PPE solution was evenly dripped onto the sponge; the control group received an equivalent volume of physiological saline. After 5 minutes of incubation in the dark, the gelatin sponge was removed, and the abdominal cavity was thoroughly rinsed three times with saline to remove residual PPE. The abdomen was then sutured layer by layer. Post-operative mice were placed on a heating pad for recovery.

### 2.3 Ultrasound Imaging Monitoring

A VINNO D700LAB small animal high-frequency ultrasound system was used to monitor the mouse abdominal aorta. Mice were anesthetized with inhaled 1-2% isoflurane and placed in a supine position. After applying coupling gel, the X10-23L probe (frequency 23 MHz, adjusted detection depth 1.5 cm) was used to locate the infrarenal aorta in the long-axis view, measuring its maximum diastolic internal diameter. All ultrasound measurements were performed by an operator blinded to the experimental groups.

### 2.4 Tissue Sample Collection

#### Samples for molecular biology experiments

Mice were anesthetized and euthanized. The vasculature was flushed via cardiac apex perfusion with ice-cold PBS. The infrarenal abdominal aorta was rapidly dissected, rinsed with ice-cold PBS, immediately snap-frozen in liquid nitrogen, and subsequently stored at −80°C.

#### Samples for morphological experiments

Mice were anesthetized and perfused via the cardiac apex sequentially with ice-cold PBS and 4% paraformaldehyde for systemic fixation. The abdominal aorta was dissected, surrounding adipose tissue was removed, and it was placed in 4% paraformaldehyde for fixation at 4°C for 24-48 hours, followed by dehydration and embedding.

### 2.5 Protein Extraction and Quantification

#### Tissue samples

Mechanically homogenized in pre-cooled RIPA lysis buffer (containing protease and phosphatase inhibitors) and lysed at 4°C for 60 min. Lysates were centrifuged at 4°C, 12,000 rpm for 15 min, and the supernatant was collected as total protein. Protein concentration was determined using the BCA method, and sample concentration was calculated by plotting a standard curve.

#### Cultured cells

Treated cells were washed with PBS 3 × 3 min. 200 µL of pre-cooled RIPA lysis buffer was added per culture dish, and cells were placed on ice for 3 min. Cells were scraped and lysed on a rotator at 4°C for 20 min. Centrifuged at 12,000 × g, 4°C for 15 min. The supernatant was collected, quantified by BCA assay, aliquoted, and stored at −80°C for Western blot.

### 2.6 Western Blot Analysis

Protein samples were separated by SDS-PAGE and transferred to PVDF membranes. Membranes were blocked with 5% skim milk and incubated with corresponding primary antibodies overnight at 4°C. The next day, after washing, membranes were incubated with HRP-conjugated species-specific secondary antibodies at room temperature for 1 hour, followed by three 10-minute washes with TBST. Bands were visualized using ECL chemiluminescence reagent. Band intensity was analyzed using ImageJ software. Statistical analysis was performed using the ratio of target protein to internal reference protein intensity.

### 2.7 Tissue Dehydration, Embedding, and Sectioning

Aortic tissues fixed in 4% paraformaldehyde were dehydrated through a graded ethanol series, cleared in xylene, and embedded in paraffin. Paraffin blocks were sectioned at 4 µm thickness, mounted onto slides, and baked at 70°C for 2 hours before use.

### 2.8 Histological Staining

#### Hematoxylin and Eosin (H&E) Staining

Sections were deparaffinized and rehydrated. Stained with hematoxylin for 6 min, washed with water, differentiated in 1% acid alcohol for ∼3 s, rinsed under running water, and blued in 50°C warm water. After brief treatment with 80% ethanol, sections were stained with eosin for 5 min, and 80% ethanol was applied for 10 s to stop staining. Dehydration through graded ethanol (90%, 95% ethanol for 10 s each; absolute ethanol I/II for 3 min each), clearing in xylene (I–III for 3 min each), mounted with neutral balsam, and imaged using a panoramic scanner for analysis.

#### Elastic Fiber Staining (EVG)

Sections were deparaffinized and rehydrated. Tissue areas were circled with an immunohistochemical pen. Fresh staining solution A (A1:A2:A3=5:2:2) was prepared and applied for 3 min, followed by water wash to stop. Solution B was added dropwise under a microscope until tissue turned brown (∼3 min), washed with water to stop, and treated with 95% ethanol for 5 s for deiodination. Fresh staining solution C (C1:C2=1:1) was applied for counterstaining for 2 min. Dehydration through graded ethanol, clearing in xylene, mounted with neutral balsam, and imaged with a panoramic tissue scanner.

#### Victoria Blue (VB) Staining

Sections were deparaffinized and rehydrated, and tissue was circled. Fresh oxidizing solution A (A1:A2=1:1) was prepared and applied for 5 min, washed with water to stop. After drying, bleaching solution B was added for 2 min, followed by a 5-min water wash. After brief treatment with 70% ethanol, staining solution C was added and incubated at room temperature in the dark for 24 hours. The next day, residual dye was removed with 70% ethanol, washed with water. After drying, staining solution D was added for nuclear counterstaining for 6 min, washed with water. Dehydration through graded ethanol (75%→85%→95% for 2 min each, absolute ethanol I/II for 3 min each), clearing in xylene (I–III for 3 min each), mounted with neutral balsam, and imaged with a panoramic tissue scanner.

### 2.9 Tissue Immunofluorescence

Sections underwent antigen retrieval with sodium citrate, blocked with 3% BSA, and incubated with primary antibodies overnight at 4°C. The next day, sections were incubated with corresponding species-specific fluorescent secondary antibodies at room temperature in the dark for 2 hours. Nuclei were counterstained with DAPI. Sections were mounted with anti-fade mounting medium. Images were observed and captured using a Leica inverted fluorescence microscope and analyzed.

### 2.10 qPCR

Total RNA was extracted from tissues using the TaKaRa MiniBEST Universal RNA Extraction Kit. After assessing purity and concentration with NanoDrop, cDNA was synthesized using the PrimeScript™ FAST RT reagent Kit with gDNA Eraser. Using cDNA as template, amplification was performed on a real-time quantitative PCR system using TB Green^®^ Premix Ex Taq II (Tli RNaseH Plus). The program: 95°C for 30 s pre-denaturation; 40 cycles of 95°C for 5 s, 60°C for 32 s. Relative gene expression was calculated using the 2^-ΔΔCt^ method.

### 2.11 Whole Transcriptome Sequencing and Analysis

Transcriptome sequencing of mouse AAA lesion and Sham group vascular tissues was commissioned to Servicebio using the DNBSEQ-T7RS high-throughput sequencing platform for PE150 paired-end sequencing. Post-sequencing data underwent quality control with fastp^26^, alignment to the reference genome using STAR^27^ to obtain gene expression quantification data. Differential expression gene analysis and functional enrichment analysis were performed using R packages DESeq2^28^ and clusterProfiler^29^, etc.

### 2.12 Re-analysis of Public Single-Cell Transcriptome Sequencing Data

To investigate the cellular source and expression characteristics of PGC-1α (gene *Ppargc1a*) in AAA, we conducted an in-depth analysis of the published single-cell RNA sequencing dataset (GSE152583)^30^. This dataset includes aortic tissue samples from mouse AAA models (7-day and 14-day elastase perfusion groups) and sham operation controls. Data analysis utilized the scanpy package^31^, strictly following standard single-cell data processing pipelines: after loading Cell Ranger-aligned and quantified data, multi-level quality control was applied to retain cells meeting the following criteria: (1) ≥200 genes detected per cell; (2) each gene expressed in at least 3 cells; and (3) mitochondrial gene percentage <10%. Additionally, cell-cycle-related variation was regressed out using cell-cycle scoring, and potential doublets were identified and removed using Scrublet. Subsequently, log-normalization, highly variable gene selection, PCA dimensionality reduction were performed. Harmony algorithm was used for batch effect correction. Finally, cell clustering was achieved based on UMAP visualization and the Leiden algorithm (resolution = 0.6). Nine cell types were identified and annotated: Vascular Smooth Muscle Cells (VSMC), Endothelial Cells (EC), Fibroblasts (Fibro), Macrophages (Mac), Monocytes (Mono), Dendritic Cells (DC), Natural Killer Cells (NK), B cells (B), and Neurons (Neuron).

### 2.13 Genotyping

1-2 mm of mouse tail tip tissue was clipped, disinfecting the tail with alcohol swabs before and after. Tissue was placed in a 1.5 mL EP tube with 200 µL of fresh lysis buffer (1 mL PBND buffer containing 20 µL proteinase K), and digested in a 55°C water bath overnight. The next day, the lysate was heated in a 100°C metal bath for 10 min, centrifuged, and placed on ice for later use. PCR reaction mix was prepared on ice: adding 2×Fast sTaq PCR Master Mix, PCR Forward Primer (10 µM), PCR Reverse Primer (10 µM), DNA template, and nuclease-free water sequentially into sterile, enzyme-free PCR tubes. PCR program: Pre-denaturation 95°C 3 min; 40 cycles: 95°C 30 s, 58°C 30 s, 72°C 1 min; final extension 72°C 7 min. Agarose gel of appropriate concentration was prepared based on expected PCR product size, placed in an electrophoresis tank with 1X TAE buffer. DNA Marker and PCR products were loaded into wells, and electrophoresis was performed at 100V for 30 min. After electrophoresis, gels were imaged using a gel imaging system, and target genotypes were screened based on band patterns.

### 2.14 Extraction, Passage, and Gene Knockout Model Operations of Mouse Primary VSMCs

Primary VSMCs were isolated from aortas of 10-12 week-old C57BL/6J wild-type mice or tamoxifen-induced (100 mg/kg for 5 consecutive days, observed for 3 days after cessation) *Ppargc1a*^flox/flox^; *Itga8-Cre^T2^* conditional knockout mice (VSMC-specific PGC-1α KO) under sterile conditions.

#### Extraction protocol

Mice were anesthetized with isoflurane and euthanized by cervical dislocation. After surface disinfection with 75% ethanol, the abdomen was opened to expose the abdominal aorta. The aorta along with the heart was completely dissected and immersed in ice-cold PBS. Under a microscope, fat, adherent tissue on the heart, and the adventitia were removed, retaining the medial segment. Two-step enzymatic digestion: (1) 0.15% collagenase II incubation at 37°C for 10 min (before stripping adventitia); (2) Minced medial tissue was placed in fresh collagenase II and digested at 37°C for another hour. Digestion was terminated by adding complete medium containing 20% FBS. Cell suspension was collected and centrifuged under conditions adjusted for cell type: Wild-type VSMCs at 1500 rpm × 3 min, PGC-1α KO VSMCs at 1000 rpm × 2 min (to reduce mechanical damage). Cells were resuspended and seeded into gelatin-coated T25 flasks. Adherent growth was observed after 48 hours. All experiments were performed within passages P1–P3.

#### Passage and plating

When cells reached 80–90% confluency, medium was discarded, and trypsin was added at 37°C until over 80% of cells became round (∼1 min), promptly neutralized with complete medium. Cells were gently pipetted to collect, centrifuged under the corresponding conditions, resuspended in complete medium with 20% FBS, and either passaged (T25) or seeded into multi-well plates as required by experiments.

### 2.15 Cellular Senescence Model Construction

When VSMCs in dishes or plates reached 80% confluency, complete medium was discarded, and cells were gently washed with PBS 2-3 times. The experimental group received serum-free medium containing 10-6 M Ang II, while the Control group received serum-free medium only. Cells were incubated at 37°C, 5% CO_2_ for 24 hours for subsequent detection.

### 2.16 CCK-8 Cell Viability Assay

Cells were seeded at a density of 1,000 – 3,000 cells per well in 6-well plates and incubated at 37°C in a 5% CO_2_ incubator for 24 hours to allow attachment. The culture medium was then aspirated, and cells were washed 2-3 times with PBS. Treatment solutions— either Ang II or Ang II+Met— were added, and cells were incubated for an additional 24 hours under the same conditions. Following treatment, the drug-containing medium was removed, and cells washed 3 times with PBS. Subsequently, 90 µL of complete medium + 10 µL of CCK-8 reagent was added to each well. Plates were incubated at 37°C in the dark, and absorbance at 450 nm (OD_450_) was measured every 30 minutes. The reaction was terminated once the OD value reached 1.0-2.0. Relative cell viability was calculated normalized to the control group.

### 2.17 Senescence-Associated β-Galactosidase (SA-β-gal) Staining

Sterile coverslips were pre-placed in 6-well plates, and VSMCs were seeded and cultured for 24 hours to allow attachment. After treatment with Ang II ± Met for 24 hours (as in 2.15), the drug-containing medium was removed, and cells were washed 3 times with PBS. Cells were then fixed with 1 mL SA-β-gal fixation solution at room temperature for 15 min, followed by three 2-min PBS washes. Fresh SA-β-gal staining working solution (containing X-gal) was added, and plates were incubated overnight at 37°C in dark, with the wells sealed with parafilm to prevent evaporation. After staining, SA-β-gal-positive cells (exhibiting blue staining) were washed 3 times with PBS, dehydrated through graded ethanol series (90% and 95% ethanol for 10 s each; anhydrous ethanol I/II for 3 min each), cleared in xylene (three changes, 3 min each), and mounted with neutral balsam. Images were captured using a panoramic tissue scanner, and the proportion of SA-β-gal-positive cells was quantified.

### 2.18 Immunofluorescence Staining

Cell processing followed the protocol in Section 2.17 (treatment with Ang II ± Met for 24 h). **Fixation:** Cells were fixed with 4% paraformaldehyde for 15 min at room temperature, followed by three 3-min PBS washes. **Permeabilization:** Cells were permeabilized with 0.1% Triton X-100 for 30 min, followed by three 3-min PBS washes. **Blocking:** Cells were blocked with 3% BSA for 30 min at room temperature, followed by three 3-min PBS washes. **Primary antibody:** Cells were incubated with primary antibody overnight at 4°C. **Secondary antibody incubation:** Fluorophore-conjugated secondary antibodies were applied and incubated for 1 h at room temperature in the dark, followed by three 3-min PBS washes. **Mounting:** Nuclei were counterstained with DAPI, and coverslips were mounted with antifade mounting medium. After final washes, coverslips were gently lifted with forceps, excess liquid was blotted away, and images were acquired using a Leica (STELLARIS 5) inverted fluorescence microscope. Fluorescence intensity was quantified with ImageJ.

### 2.19 Cell Scratch Migration Assay

VSMCs were seeded in 6-well plates and cultured for 24 hours to allow attachment. After 24 h treatment with Ang II ± Met, the drug-containing medium was removed, cells were washed 3 times with PBS, and fresh complete culture medium was added. A sterile 1 mL pipette tip was used to create a straight scratch wound perpendicular to the well bottom. Images were captured at 0 hours (inverted microscope, ×10 objective), and cells were further cultured with time-lapse imaging at 12 and 24 hours. Scratch closure area was measured using ImageJ, and cell migration rate was calculated.

### 2.20 Transmission Electron Microscopy Observation of Mitochondrial Ultrastructure

Treated cells were collected, resuspended and thoroughly mixed in electron microscopy (EM) fixative, and fixed at 4°C for 2-4 hours. After fixation, cells were washed 3 times (3 min each) with 0.1 M PB buffer (pH=7.4) and centrifuged to obtain a pellet.

A 1% agarose solution was prepared in advance by heating and dissolving in water, then slightly cooled. The agarose was added to the microcentrifuge tube containing the cell pellet; before the agarose solidified, the pellet was carefully lifted with fine forceps and fully embedded within the agarose.

#### Post-fixation

Samples were fixed in 1% osmium tetroxide prepared with 0.1 M phosphate buffer (pH=7.4) at room temperature in the dark for 2 hours. After post-fixation, samples were rinsed three times (15 min each) with 0.1 M PB (pH=7.4).

#### Dehydration

Graded ethanol series: 30%→50%→70%→80%→95%→100% ethanol I→100% ethanol II (20 min per step), followed by two 15-min rinses in 100% acetone.

#### Infiltration and embedding

Samples were performed sequentially with acetone: Embed-812 resin mixtures: 1:1 (v/v, 37°C, 2-4 h) → 1:2 (v/v, 37°C, overnight) → pure Embed-812 resin (37°C, 5-8 h). Finally, samples were transferred into embedding molds filled with fresh pure Embed-812 resin, oriented appropriately, and polymerized at 37°C overnight, followed by curing at 60°C for 48 hours to form resin blocks.

#### Ultramicrotomy and staining

Ultrathin sections (60-80 nm) were cut and collected onto 150-mesh copper grids. Sections were stained sequentially as follows: (1) Uranyl acetate staining: 2% uranyl acetate in saturated ethanol, 8 min in the dark; rinsed 3 times alternating with 70% ethanol and ultrapure water. (2) Lead citrate staining: 2.6% lead citrate, 8 min (protected from CO2); rinsed 3 times with ultrapure water and gently blotted with filter paper. Grids were air-dried overnight at room temperature and examined under a transmission electron microscope. Mitochondrial morphology was assessed with emphasis on cristae integrity, swelling, and vacuolization.

### 2.21 Intracellular ATP Assay

After cell lysis, samples were sonicated for ∼5 min, heated in a boiling water bath for 2 min, and rapidly cooled under running water. The lysates were then centrifuged at 10,000 *g* for 10-15 min at 4°C, and the supernatants were collected and kept on ice. A stock ATP standard (200 µM) was serially diluted to generate a standard curve. The ATP detection enzyme and ATP assay buffer were mixed at a ratio of 1:50 to prepare the working reagent. For each well of a 6-well plate, 100 µL working solution was added and equilibrated for 5 min at room temperature. Subsequently, 20 µL of standard solution or prepared sample was added to the respective wells, mixed thoroughly by gentle crosswise shaking, and luminescence signals were measured immediately using a microplate reader in chemiluminescence mode. ATP concentrations were calculated based on the standard curve.

### 2.22 Intracellular ROS Measurement

Cells were seeded on coverslips following the protocol in Section 2.18. After 24 hours treatment with Ang II ± Ang II+Met, cells were washed 3 times with PBS and incubated with freshly prepared H_2_DCFDA (10 μM) at 37°C in 5% CO2 for 30 min in the dark. Following incubation, cells were washed 3 × 3 min with PBS, counterstained with DAPI, and mounted. Fluorescence images were acquired using a Leica inverted fluorescence microscope, and intracellular ROS levels were quantified by measuring mean fluorescence intensity (ImageJ).

### 2.23 Statistical Analysis

Data are presented as mean ± standard deviation (SD). Statistical analyses were performed using GraphPad Prism 9.0. Prior to hypothesis testing, normality (e.g., Shapiro – Wilk test) and homogeneity of variances (e.g., Brown – Forsythe test) were assessed. For data meeting the assumptions of normality and equal variance: (1) comparisons between two independent groups were performed using unpaired, two-tailed Student’s t-test; (2) comparisons among ≥3 groups were conducted by one-way analysis of variance (ANOVA), followed by Tukey’ s test (equal group sizes) or Tukey– Kramer test (unequal group sizes) for post hoc pairwise comparisons. Experiments with two independent variables were analyzed by two-way ANOVA. For data violating parametric assumptions: Mann–Whitney U test (two groups) or Kruskal–Wallis test with Dunn’s post hoc test (≥3 groups) were applied. A P value < 0.05 was considered statistically significant.

## 3. Results

### 3.1 Exacerbated Cellular Senescence and Transcriptional Abnormalities During Mouse AAA Progression

Cellular senescence is an irreversible state of cell cycle arrest, an inevitable physiological process for cells and the entire organism. Excessive cellular senescence is regarded as a key driver or contributing factor in the pathogenesis of multiple diseases^17,19,32^. To explore whether cellular senescence exists and its degree during AAA progression, we constructed a mouse AAA model using local PPE application^25,33^. Fourteen days after model induction, Western blot results of lesion vascular tissues showed that the expression levels of senescence marker molecules p53 and p21 were significantly increased in the Model group compared to the Sham group (Figure 1A-1C). Immunofluorescence further confirmed that the proportions of p53-positive and p21-positive cells within the abdominal aortic tissue of Model mice were significantly increased (Figure 1D-1G). Histologically, H&E staining showed enlarged interstitial spaces in the aortic wall and perivascular tissues in the Model group, with vacuolar degeneration of medial SMCs and inflammatory cell infiltration (Figure 1H). EVG and VB staining results indicated significant elastic fiber fragmentation in the aortic media of the Model group (Figure 1H).

**Figure 1.**
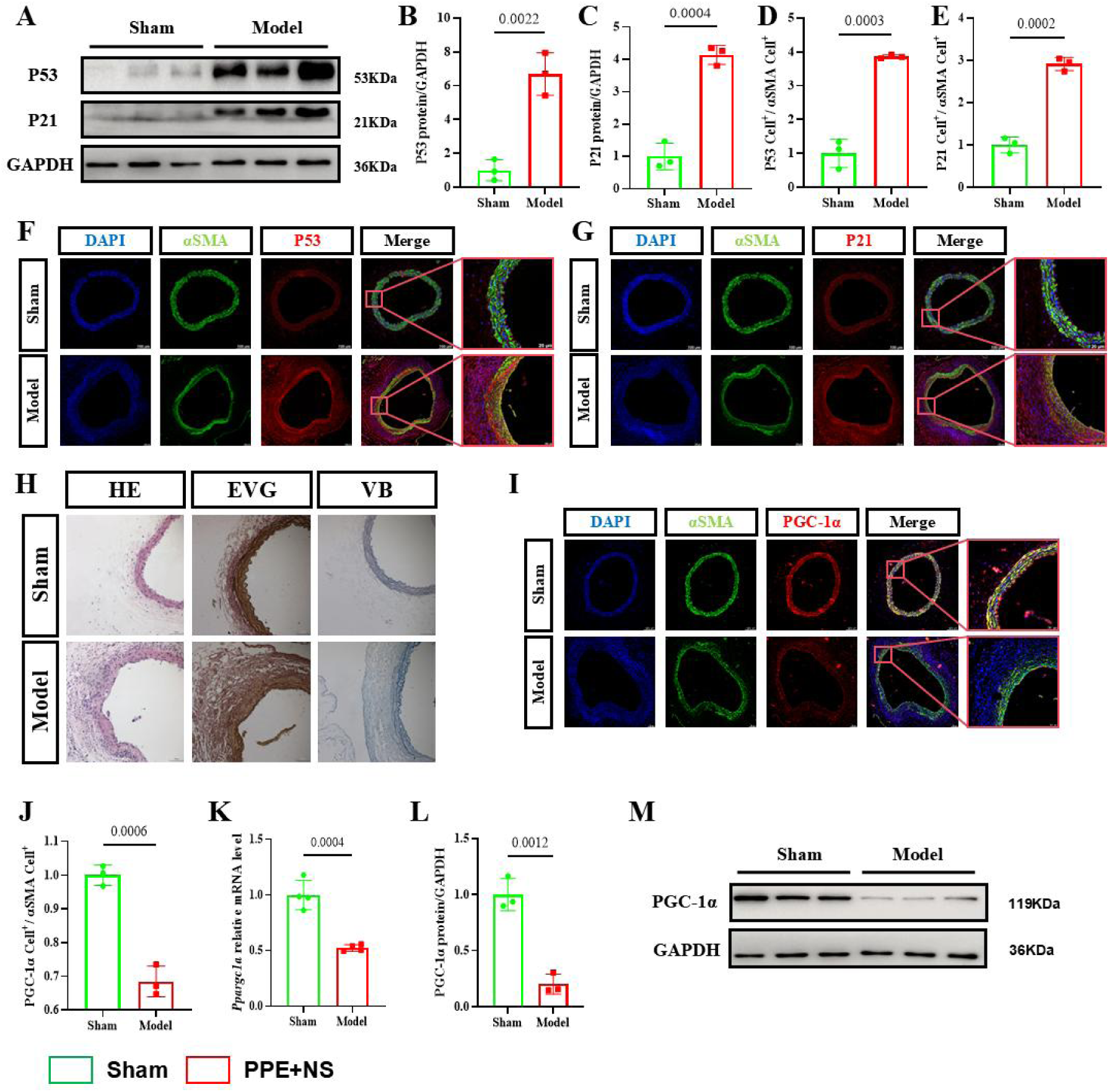
Cellular senescence is exacerbated and accompanied by transcriptional dysregulation during abdominal aortic aneurysm (AAA) progression in mice. (A) Western blot analysis of p53 and p21 protein expression in abdominal aortic tissues from control (Sham) and AAA mice. (B, C) Quantification of band intensities from (A) (n = 3 mice/group). (D, E) Quantification of p53- and p21-positive cells, respectively, from immunofluorescence staining in (F) and (G) (n = 3). (F, G) Representative immunofluorescence images showing the percentage of p53-and p21-positive cells in abdominal aortic sections from Sham and AAA mice. (H) Hematoxylin–eosin (H&E), Elastin–van Gieson (EVG), and Victoria Blue (VB) staining illustrating pathological alterations in aortic tissues (n = 3). (I) Immunofluorescence detection of PGC-1α (peroxisome proliferator-activated receptor gamma coactivator 1-alpha)-positive cells in aortic sections. (J) Quantification of PGC-1α-positive cells from (I) (n = 3). (K) qRT-PCR analysis of *Ppargc1a* mRNA expression in aortic tissues (n = 4). (L) Quantification of PGC-1α protein levels from Western blot in (M) (n = 3). (M) Western blot of PGC-1α in aortic lysates. Data are presented as mean ± SD. Statistical significance was assessed by two-tailed unpaired Student’s *t*-test (B, C, D, E, J, K, L); *p* < 0.05 was considered statistically significant.

To investigate the cause of increased senescence in AAA tissue, we performed whole transcriptome RNA sequencing analysis on Model group AAA tissues and Sham group normal abdominal aortic tissues to screen for differentially expressed genes. Gene Set Enrichment Analysis (GSEA) indicated that GO terms significantly enriched among downregulated genes, including *Ppargc1a* (PGC-1α protein gene), were related to lipid and carbohydrate metabolism and regulation, such as *circadian rhythm*, *steroid metabolic process*, *regulation of lipid metabolic process*, *hexose biosynthetic process*, *fatty acid oxidation*, *lipid modification*, *lipid oxidation*, *positive regulation of lipid metabolic process*, *regulation of fatty acid oxidation*, *monosaccharide biosynthetic process*, *positive regulation of lipid biosynthetic process*, and *regulation of lipid biosynthetic process* (Figure S1A-S1D, Figure S2A-S2D, Figure S3A-S3D). Furthermore, KEGG pathways significantly enriched with genes including *Ppargc1a* were: *AMPK signaling pathway*, *thermogenesis*, and *Huntington disease* (Figure S4A-S4C). These results collectively suggest *Ppargc1a* primarily participates in cellular energy metabolism-related processes. Western blot results showed that both protein and mRNA levels of PGC-1α were significantly reduced in the AAA of the Model group (Figure 1K-1M).

To further clarify the cellular source of PGC-1α, we integrated public single-cell RNA sequencing data (GSE152583) for cell subpopulation annotation of mouse abdominal aorta (Figure S5A). Analysis showed that *Ppargc1a* was mainly expressed in VSMCs, with relatively low expression levels in other cell types (Figure S5B). This result was validated by double immunofluorescence staining (α-SMA⁺/PGC-1α⁺): PGC-1α⁺ signal was predominantly localized to the medial layer of VSMCs, and its expression intensity trend was consistent with Western blot results (Figure 1I-1J), confirming VSMCs as the key target cells for PGC-1α dysfunction in AAA.

### 3.2 Low-Dose Metformin Effectively Ameliorates Cellular Senescence in AAA Tissue

Clinical evidence indicates that metformin attenuates aneurysm expansion and lowers the risk of AAA rupture in patients^7,8^. Emerging evidence further suggests that metformin exerts pleiotropic anti-senescence effects^10,34^, prompting us to hypothesize that it may attenuate AAA progression, at least in part, via suppression of cellular senescence. To test this, we administered metformin at 0 (vehicle), 50, or 150 mg/kg/day by oral gavage to mice with PPE-induced AAA for 14 days. Both 50 mg/kg and 150 mg/kg/day significantly inhibited aneurysm expansion (Figure S6A-S6B); notably, the low-dose regimen (50 mg/kg/day) more potently suppressed the expression of senescence markers p53 and p21 in AAA tissue (Figure S7A-S7C). This dose corresponds to the clinically relevant range (∼ 1/10 – 1/20 of the LD_50_ in C57BL/6J mice) and did not cause hypoglycemia during treatment (Figure S7D), and was thus selected for subsequent experiments.

In an independent cohort, AAA mice treated with 50 mg/kg/day metformin for 14 days exhibited significantly reduced aneurysm diameters, as assessed by both gross morphological and ultrasound imaging, compared with untreated AAA controls (Figure 2A-D). Consistent with this, both Western blot and qRT-PCR analyses confirmed that metformin robustly suppressed p53 and p21 expression at both the protein and transcript levels (Figure 2E-2I). Immunofluorescence staining further demonstrated a marked reduction in the proportion of p53^+^ and p21^+^ cells in the aortic wall (Figure 2J-2M), collectively establishing metformin’s anti-senescence effect in vivo.

**Figure 2.**
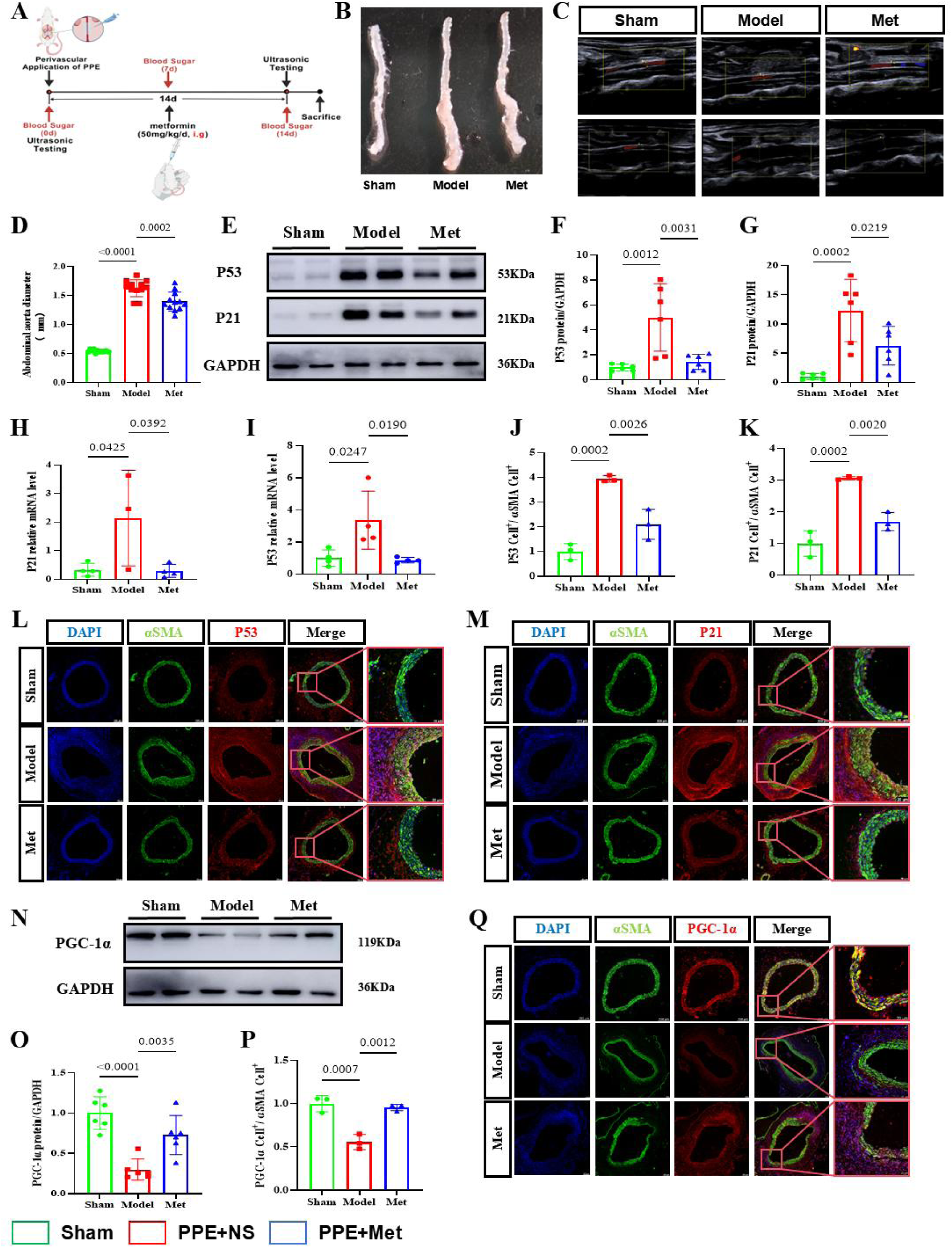
Metformin alleviates cellular senescence in AAA tissues. (A) Experimental timeline: 10–12-week-old C57BL/6J mice underwent baseline ultrasound measurement of abdominal aortic diameter, followed by AAA induction via periaortic application of porcine pancreatic elastase (PPE). After 14 days of recovery, mice received daily intragastric (i.g.) administration of metformin (50 mg/kg) or vehicle (sterile ultrapure water) for 14 days. Final aortic diameter was assessed by ultrasound before sacrifice. (B) Representative gross morphology of excised aortas (left to right: Sham, Model, Metformin). (C) Representative ultrasound images of abdominal aortas pre- and post-modeling. (D) Quantification of maximal aortic diameter from (C) (n = 12/group). (E) Western blot of p53 and p21 in aortic lysates from Sham, Model, and Metformin groups. (F, G) Densitometric quantification of (E) (n = 6). (H, I) qRT-PCR analysis of *Trp53* and *Cdkn1a* (*p21*) mRNA levels (n = 3∼4). (J, K) Quantification of p53- and p21-positive cells from immunofluorescence in (L) and (M) (n = 3). (L, M) Immunofluorescence staining for p53 and p21, respectively. (N) Western blot of PGC-1α. (O) Quantification of (N) (n = 6). (P) Quantification of PGC-1α-positive cells from (Q) (n = 3). (Q) Immunofluorescence staining for PGC-1α. Data are mean ± SD. One-way ANOVA with Tukey’s post hoc test was used (D, F, G, H, I, J, K, O, P); *p* < 0.05 was considered significant.

To probe the underlying mechanism, we examined PGC-1α expression. Metformin treatment markedly restored PGC-1α protein levels in mouse aortic tissue (Figure 2N-2O) and increased the fraction of PGC-1α^+^ cells, as evidenced by immunofluorescence (Figure 2P-2Q). These findings collectively support a mechanistic link between metformin, PGC-1α upregulation, and alleviation of vascular senescence in AAA ^35^.

### 3.3 Metformin Reverses VSMC Phenotypic Switching During AAA Progression

VSMCs exhibit high phenotypic plasticity and, under specific pathological conditions, can transition from a contractile phenotype to a secretory, synthetic, or other dedifferentiated states ^36^, thereby losing their fundamental capacity to maintain vascular wall structural integrity and tone regulation. Accumulating evidence indicates that a chronic pro-inflammatory microenvironment serves as a pivotal driver of VSMC phenotypic switching ^37–39^. To determine whether such an inflammatory milieu exists during AAA progression, we assessed the expression levels of pro-inflammatory cytokines IL-6 and TNF-α in aortic tissues. Western blot analysis revealed significantly elevated IL-6 and TNF-α protein levels in the Model group (P < 0.01), whereas metformin treatment dose-dependently suppressed their expression (Fig. 3A – C). Immunofluorescence staining further demonstrated markedly enhanced IL-6 and TNF-α signals in the medial layer of the aorta in the Model group (Fig. 3D, E, G, H), indicating sustained local inflammatory activation—thereby establishing the necessary pathological precondition for VSMC phenotypic switching. Notably, metformin intervention effectively attenuated this pro-inflammatory trend.

**Figure 3.**
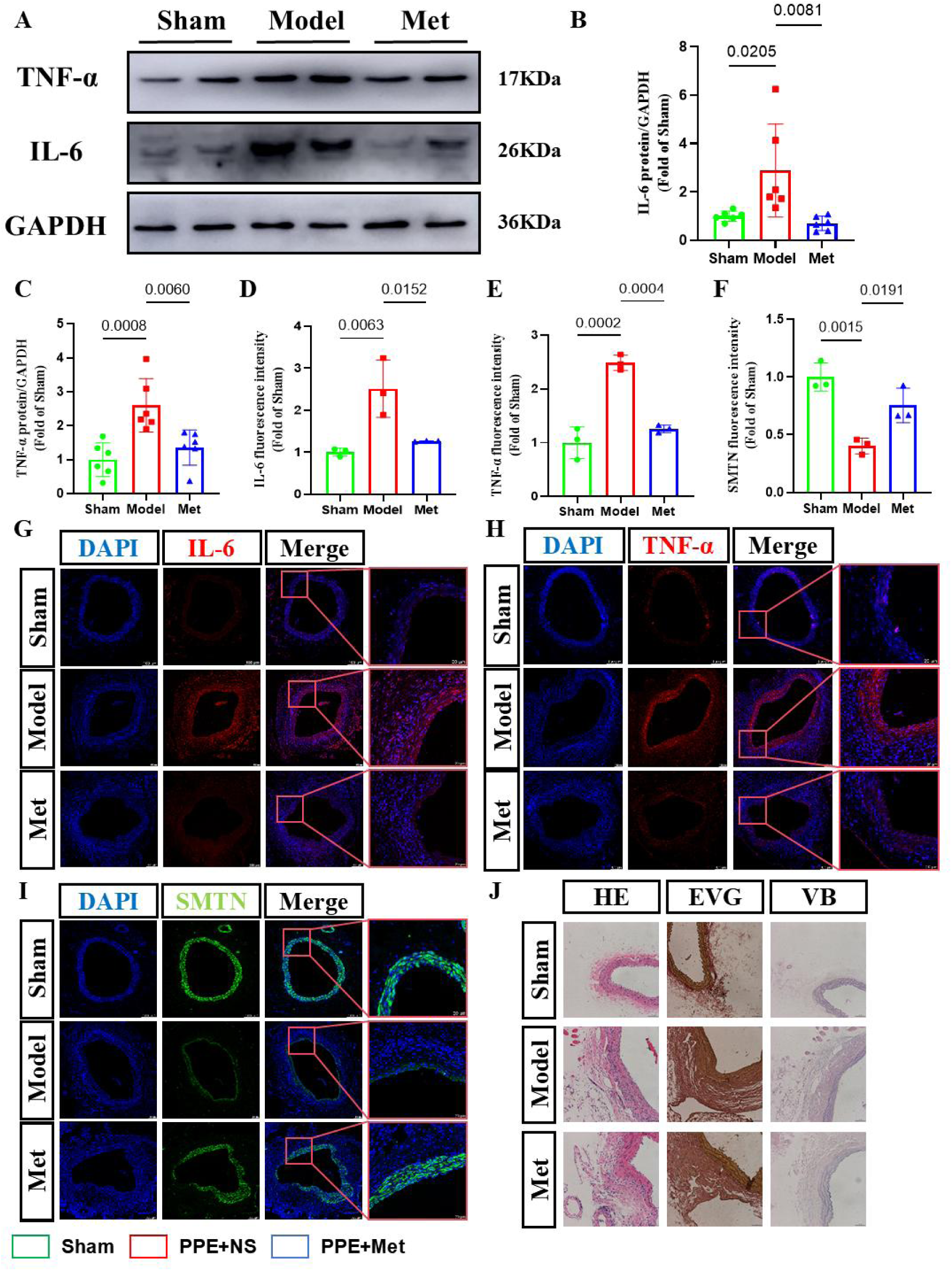
Metformin reverses vascular smooth muscle cell (VSMC) phenotypic switching during AAA progression. (A) Western blot of TNF-α and IL-6 in aortic tissues. (B, C) Quantification of (A) (n = 6). (D–F) Quantification of fluorescence intensity for TNF-α, IL-6, and smoothelin (SMTN) from (G–I), respectively (n = 3). (G–I) Immunofluorescence staining for TNF-α, IL-6, and SMTN in aortic sections. (J) H&E, EVG, and VB staining of aortic pathology (n = 3). Data are mean ± SD. One-way ANOVA with Tukey’s post hoc test (B, C, D, E, F); *p* < 0.05 was considered significant.

Building on this, we further evaluated whether VSMCs indeed undergo a phenotypic shift from contractile to secretory/synthetic states. Smoothelin (SMTN), a core component of the VSMC contractile apparatus, is widely recognized as a critical molecular marker for the maintenance of the contractile phenotype; its downregulation is a well-established indicator of VSMC transition toward a synthetic/secretory phenotype ^40^. Immunofluorescence analysis showed a significantly reduction in SMTN signal intensity in the medial layer of the Model group (P < 0.001), whereas metformin treatment robustly restored SMTN expression (Figure 3F, 3I), providing direct evidence for the loss of the contractile phenotype and acquisition of a secretory phenotype in VSMCs. Histological analyses further corroborated this conclusion: Hematoxylin – eosin (H&E) staining revealed severe medial disorganization and pronounced inflammatory cell infiltration in the Model group, whereas metformin markedly alleviated immune cell infiltration, improved medial structural integrity, and reduced the adventitial – medial separation (Figure 3J). Additionally, EVG and VB staining demonstrated reduced elastin fragmentation and more orderly elastic fiber arrangement in the metformin-treated group (Figure 3J), collectively suggesting that metformin preserves vascular wall architecture and functional homeostasis—likely by reversing VSMC phenotypic switching.

### 3.4 Metformin Suppresss Senescence in Primary VSMC and Rescues PGC-1α Downregulation

To validate metformin’s anti-senescence in VSMCs, we isolated and cultured primary VSMCs from mouse aortas (Figure S8A) and induced senescence using angiotensin II (Ang II), with or without metformin co-treatment. Based on CCK-8 cell viability assays, we selected 1 mM metformin as the optimal concentration—sufficient to significantly ameliorate Ang II – induced cell viability loss without cytotoxicity (Figure S8B). Subsequent analyses revealed that metformin (1 mM) markedly attenuated Ang II–driven upregulation of senescence markers p53 and p21 at the protein level (Figure 4A-4C). Consistently, SA-β-gal staining demonstrated a substantial reduction in SA-β-gal⁺ cells upon metformin treatment (Figure 4F-4G), confirming its ability to delay VSMC senescence. Moreover, wound-healing assays showed that metformin partially restored the impaired migratory and proliferative capacity of Ang II–treated VSMCs, as evidenced by accelerated wound closure (Figure 4H-4I).

**Figure 4.**
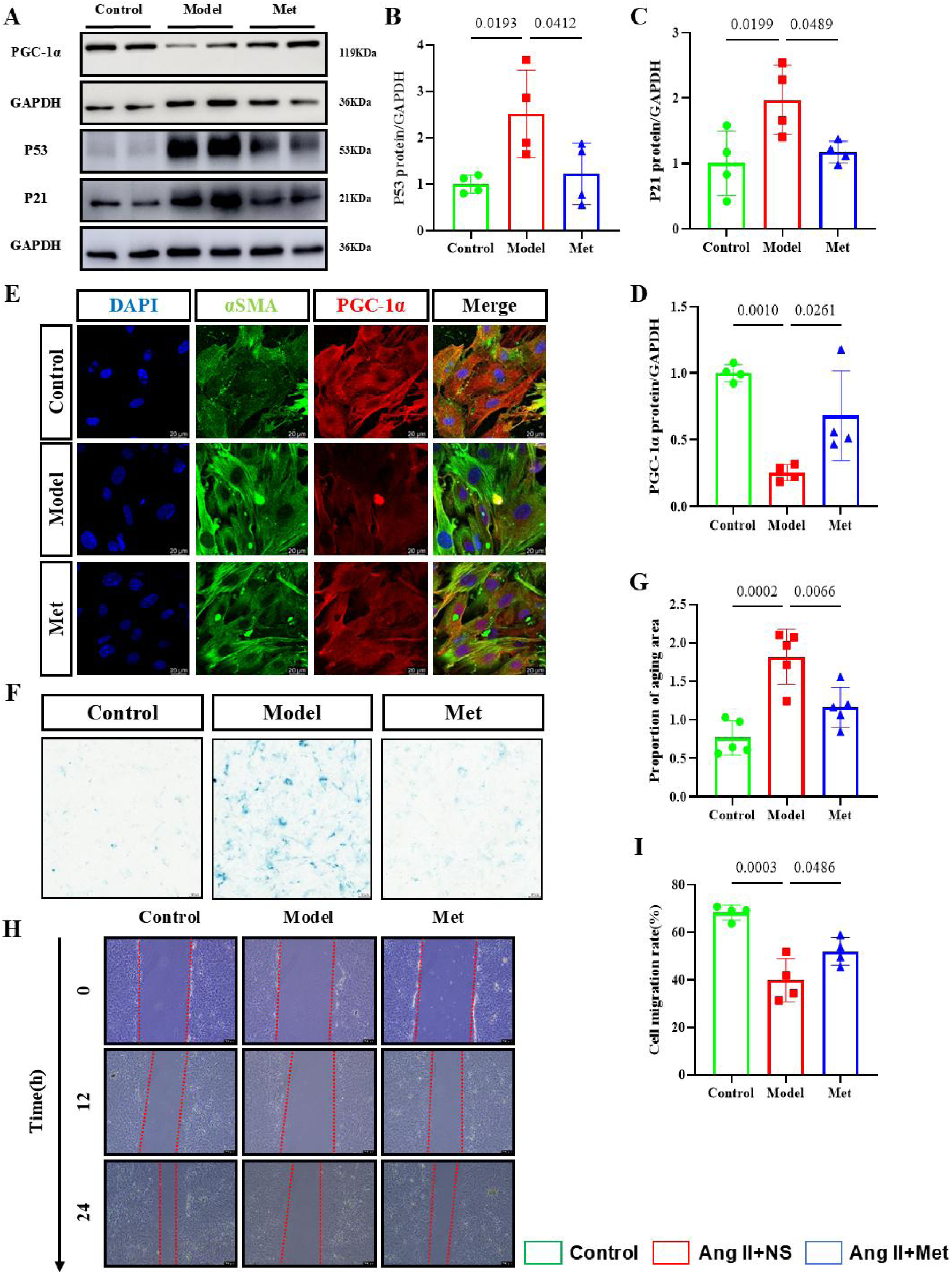
Metformin suppresses senescence and restores PGC-1α expression in primary VSMCs. (A) Western blot of p21, p53, and PGC-1α in primary VSMCs under control, PPE-induced senescence (Model), and metformin-treated (Met) conditions. (B–D) Densitometric quantification of (A) (n = 4). (E) Immunofluorescence detection of PGC-1α in VSMCs. (F) Senescence-associated β-galactosidase (SA-β-gal) staining. (G) Quantification of SA-β-gal–positive area from (F) (n = 5). (H) Wound-healing assay at 0, 12, and 24 h post-scratch. (I) Quantification of wound closure rate from (H) (n = 4). Data are mean ± SD. One-way ANOVA with Tukey’s post hoc test (B, C, D, G, I); *p* < 0.05 was considered significant.

Notably, Ang II challenge led to pronounced downregulation of *Ppargc1a* at both mRNA and protein levels; metformin treatment effectively reversed this suppression (Figure 4A, 4D-4E). These findings suggest that metformin’s anti-senescence action in VSMCs may be, at least in part, mediated through restoration of PGC-1α expression.

### 3.5 VSMC-Specific PGC-1α Deficiency Exacerbates Vascular Senescence and Abrogates Metformin’s Anti-Senescence Efficacy

To determine whether metformin attenuates AAA progression by supressing VSMC senescence in PGC-1α-dependent manner, we generated VSMC-specific *Ppargc1a* knockout mice (*Ppargc1a*^VSMC-KO^) by crossing *Ppargc1a*^flox/flox^ mice with *Itga8-Cre^T^*^2^ mice ^41^, followed by tamoxifen-induced Cre recombination. Given the highly specific expression of *Itga8* in VSMCs, *Itga8*-*CreER^T^*^2^-mediated recombination enables high cell-type specificity, and stability, and temporal control of gene manipulation ^41^. Lineage tracing using using *Rosa26*mTmG reporter mice^42^ further confirmed VSMC-restricted recombination: as eGFP^+^ signal was was exclusively detected in arterial wall VSMCs without off-target activity (Figure S9A-S9B), confirming the high efficiency and specificity of this system in VSMCs.

Ten days after completing a 5 day tamoxifen regimen, we induced AAA in *Ppargc1a*^VSMC-KO^ mice (Figure 5A, S10A-S10C). The maximal AAA diameter was significantly increased in knockout mice compared with controls. Critically, metformin failed to suppress aneurysm expansion in *Ppargc1a*^VSMC-KO^ mice (Figure 5B-5D). Western blot combined with immunofluorescence analysis confirmed that PGC-1α protein expression was significantly downregulated, and its immunofluorescence signal was nearly absent in VSMCs (Figure 5E-5H).

**Figure 5.**
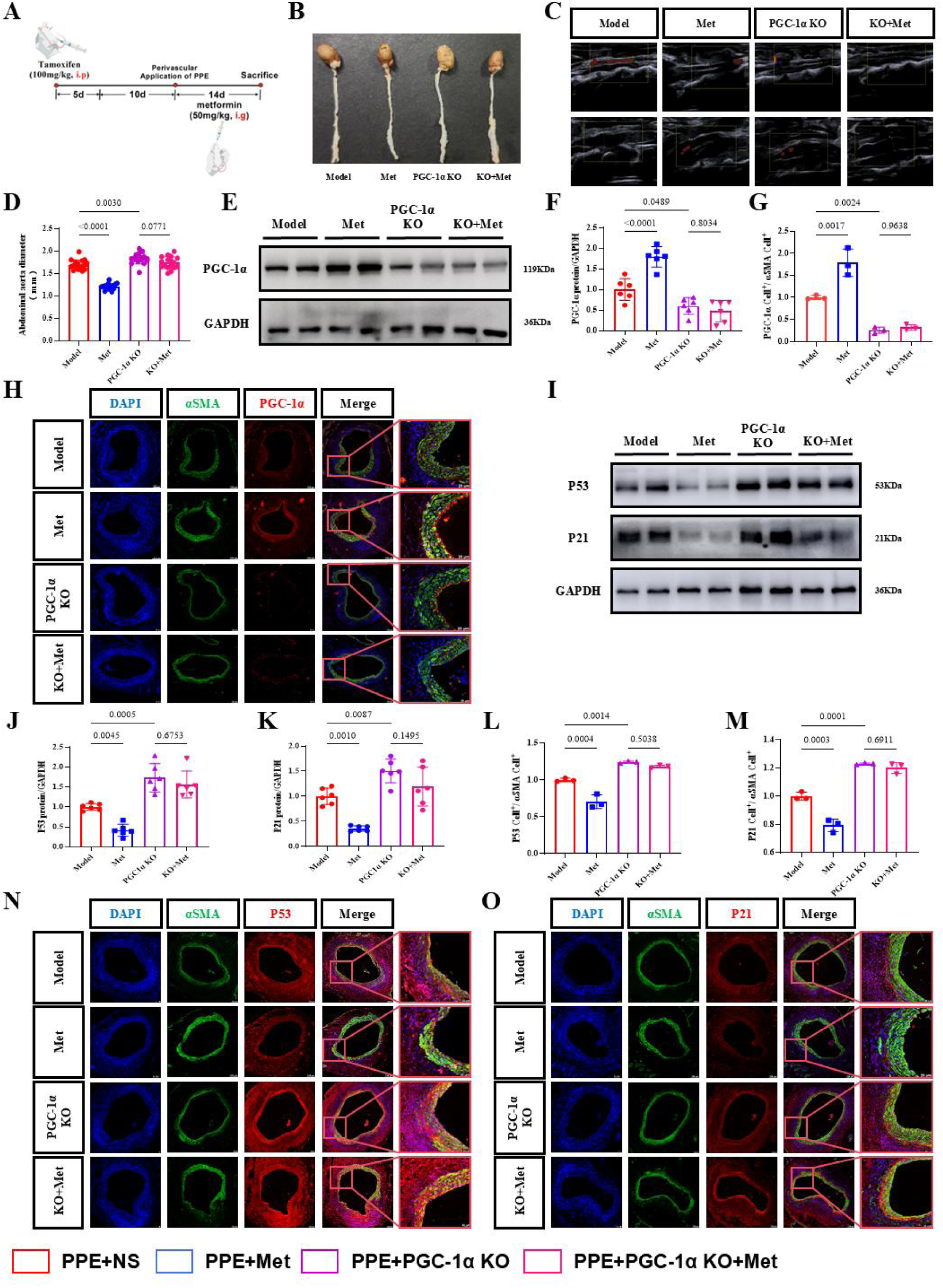
VSMC-specific *Ppargc1a* deletion exacerbates senescence and blunts the anti-senescent effect of metformin. (A) Schematic of *Ppargc1a*VSMC-KO mouse generation and AAA modeling: 10-week-old *Ppargc1a*flox/flox; *Itga8-Cre*+/− mice received intraperitoneal (i.p.) tamoxifen (100 mg/kg/day for 5 days) or vehicle (corn oil), followed by 10-day recovery and PPE-induced AAA. Metformin (50 mg/kg, i.g.) or vehicle was administered for 14 days post-modeling. (B) Representative gross aortic morphology (left to right: Model, Met, *Ppargc1a* KO, KO+Met). (C) Ultrasound images pre- and post-modeling. (D) Quantification of aortic diameter from (C) (n = 17). (E) Western blot of PGC-1α. (F) Quantification of (E) (n = 6). (G) Quantification of PGC-1α-positive cells from (H) (n = 3). (H) Immunofluorescence of PGC-1α. (I) Western blot of p53 and p21. (J, K) Quantification of (I) (n = 6). (L, M) Quantification of p53- and p21-positive cells from (N, O) (n = 3). (N, O) Immunofluorescence staining for p53 and p21. Data are mean ± SD. One-way ANOVA with Tukey’s post hoc test (D, F, G, J, K, L, M); *p* < 0.05 was considered significant.

We next assessed the impact of *Ppargc1a* deletion on cellular senescence. P53 and p21 protein levels were markedly elevated in *Ppargc1a*^VSMC-KO^ aortas, and metformin’s ability to suppress their expression was abolished in the knockout background (Figure 6I-6K). Immunofluorescence quantification further showed that PGC-1α deficiency significantly increased the proportion of P53⁺ and p21⁺ VSMCs, and metformin treatment failed to mitigate this increase (Figure 6L-6O). Together, these findings demonstrate that VSMC-specific PGC-1α loss not only exacerbates vascular senescence but also abrogates metformin’s protective effects, establishing PGC-1α as an essential downstream mediator of metformin’s anti-senescence action.

**Figure 6.**
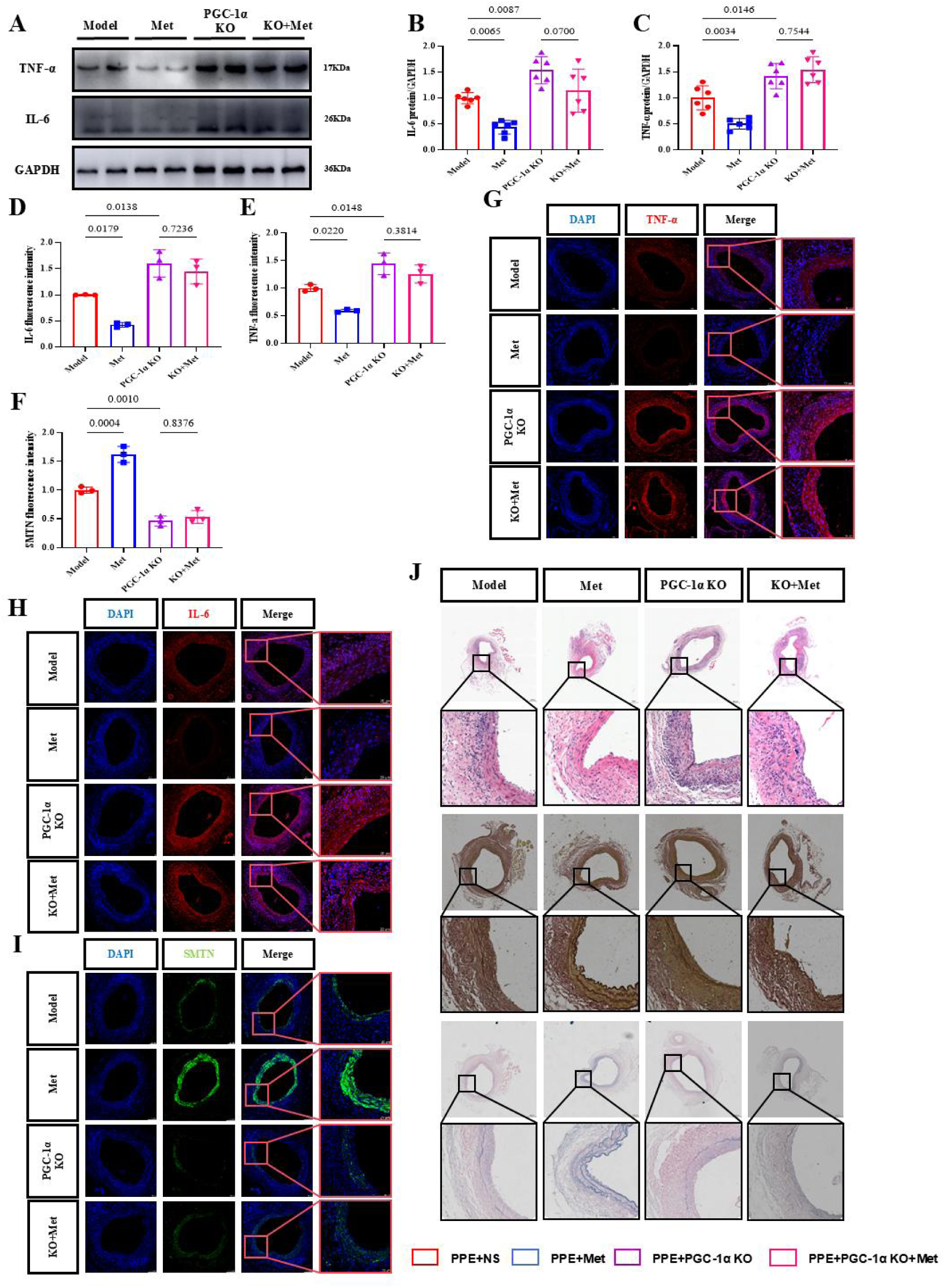
*Ppargc1a* deletion in VSMCs aggravates phenotypic switching and abrogates the therapeutic benefit of metformin. (A) Western blot of TNF-α and IL-6. (B, C) Quantification of (A) (n = 6). (D–F) Fluorescence intensity quantification for TNF-α, IL-6, and SMTN from (G–I), respectively (n = 3). (G–I) Immunofluorescence staining for TNF-α, IL-6, and SMTN. (J) H&E, EVG, and VB staining of aortic tissues (n = 3). Data are mean ± SD. One-way ANOVA with Tukey’s post hoc test (B, C, D, E, F); *p* < 0.05 was considered significant.

### 3.6 VSMC-specific PGC-1α Deficiency Exacerbates Phenotypic Modulation and Abrogates Metformin’s Vascular Protection in AAA

Cell senescence is a well-established driver of VSMC phenotypic modulation. We therefore hypothesized that PGC-1α deficiency – exacerbated senescence may further promote VSMC transition toward a pathological phenotype. Western blot analysis revealed that *Ppargc1a* deletion markedly upregulated the pro-inflammatory cytokines IL-6 and TNF-α in aortic tissues — and critically, metformin failed to attenuate this inflammatory upregulation in *Ppargc1a*^VSMC-KO^ mice (Figure 6A-6C). Consistently, immunofluorescence demonstrated intensified IL-6 and TNF-α signals within the medial layer, suggestive of expanded “secretory” VSMC populations; this effect was unresponsive to metformin treatment (Figure 6D-E, G-H).

We further examined the contractile marker smoothelin (SMTN) (encoded by *Smtn* in mice). Immunofluorescence showed that PGC-1α deficiency significantly reduced both smoothelin expression and the fraction of smoothelin^+^ VSMCs, and metformin intervention did not rescue this loss (Figure 6F-6I). Histologically, H&E staining revealed exacerbated immune cell infiltration in *Ppargc1a*^VSMC-KO^ aortas, and metformin no longer suppressed this response (Fig. 6J). Similarly, EVG and VB staining demonstrated more severe elastic fiber fragmentation in knockout mice, with complete abrogation of metformin’s protective effect (Fig. 6J).

Together, VSMC-specific *Ppargc1a* deletion abolished metformin’s ability to suppress phenotypic modulation, inflammatory cytokine release, and immune cell recruitment—collectively establishing PGC-1α as an essential mediator of metformin’s vascular protective effects in this context.

### 3.7 VSMC-Specific Ppargc1a Deficiency Disrupts Mitochondrial Struncture and Function Damage in Primary VSMCs

RNA-seq of AAA tissues from *Ppargc1a*^VSMC-KO^ mice compared to the Sham group identified 3,703 differentially expressed genes (DEGs). Intersection with MitoCarta3.0 mitochondrial compendium yielded 193 mitochondria-associated DEGs (Figure 7A). GO enrichment analysis showed significant overrepresentation in mitochondrial processes, including *mitochondrion organization*, *mitochondrial depolarization*, *mitochondrial matrix*, and *mitochondrial outer membrane* (Figure 7B-7D), while KEGG analysis implicated dysregulated amino acid metabolism (Figure 7F), collectively suggesting profound mitochondrial dysregulation in AAA tissue.

**Figure 7.**
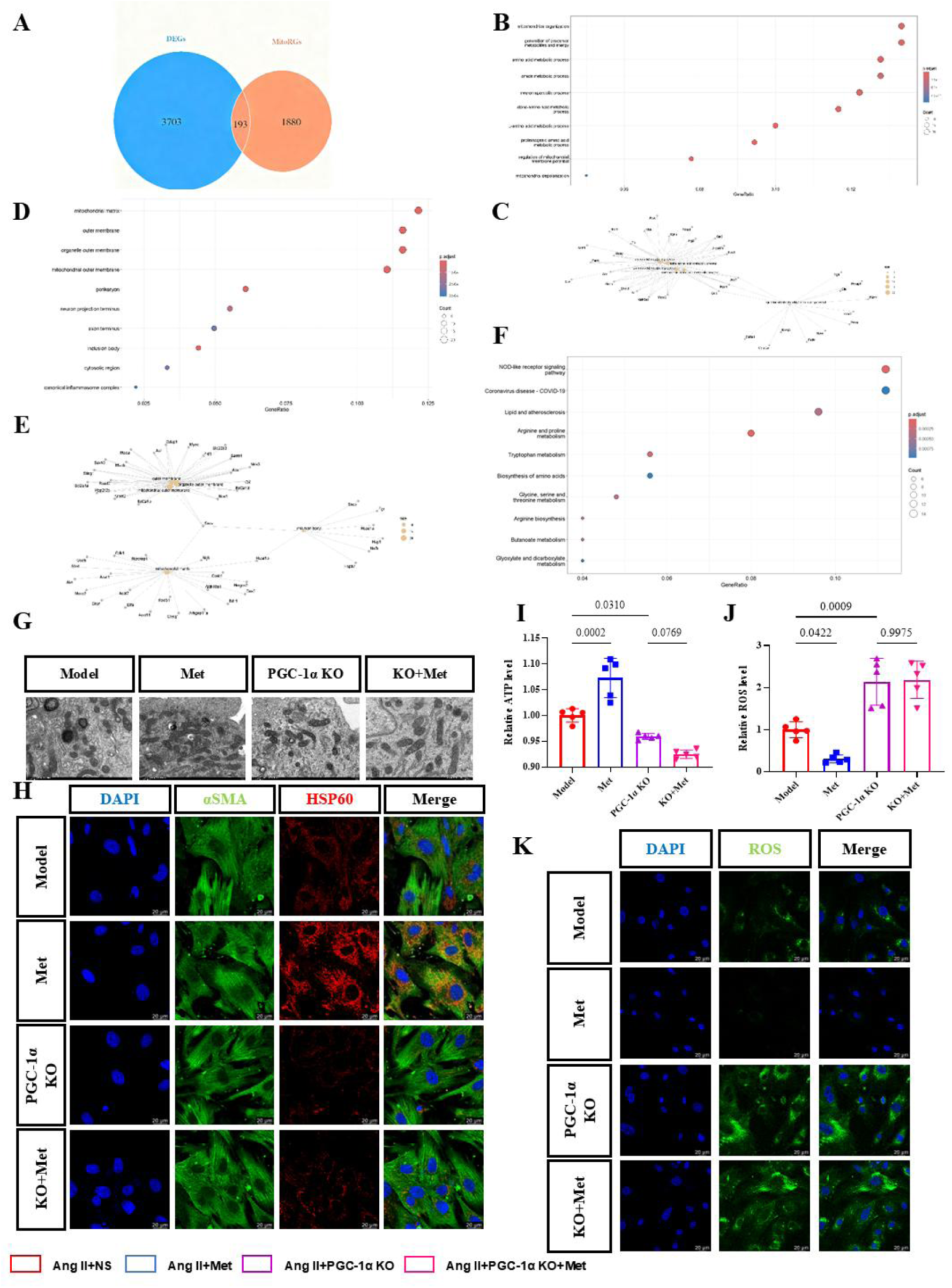
*Ppargc1a* deficiency disrupts mitochondrial morphology and function. (A) Venn diagram showing overlap between differentially expressed genes (DEGs) in *Ppargc1a* KO vs. Sham aortas and mitochondrial-related genes from public databases. (B–E) Gene Ontology (GO) enrichment analysis of the overlapping DEGs. (F) Kyoto Encyclopedia of Genes and Genomes (KEGG) pathway enrichment of overlapping DEGs. (G) Transmission electron microscopy (TEM) of primary WT and *Ppargc1a*-KO VSMCs. (H) Immunofluorescence of mitochondrial chaperone HSP60 in aortic sections. (I) Relative ATP levels in primary VSMCs (n = 5). (J) Quantification of ROS fluorescence intensity from (K) (n = 5). (K) ROS detection by H_2_DCFDA fluorescence probe. Data are mean ± SD. One-way ANOVA with Tukey’s post hoc test (I, J); *p* < 0.05 was considered significant.

To validate this at the cellular level, we isolated primary VSMCs from Model, Met (Model + Met), PGC-1α KO (Model + *Ppargc1a*^VSMC-KO^), and KO + Met (Model + *Ppargc1a*^VSMC-KO^ + Met) groups (Figure S11) and subjected them to Ang II-induced senescence. Transmission electron microscopy revealed that Model VSMCs exhibited mitochondrial matrix swelling, cristae shortening and disorganization; these ultrastructural defects were markedly ameliorated by metformin (Met group). In contrast, KO VSMCs displayed severe mitochondrial degeneration — nearly all mitochondria showed pronounced swelling and cristae shortening; with occasional autophagic engulfment forming myelin figures; metformin failed to rescue this phenotypes (Figure 7G). Consistently, immunofluorescence staining for HSP60 revealed that in Ang II – induced senescent VSMCs (Model group), mitochondrial networks were partially preserved but reduced in density; metformin treatment significantly restored mitochondrial abundance and reticular connectivity. In contrast, *Ppargc1a*-deficient VSMCs exhibited profound mitochondrial fragmentation and sparsity, with no recovery observed upon metformin intervention (Figure 7H).

Functionally, Ang II reduced cellular ATP, which metformin restored in controls but not in KO cellst (Figure 7I). Similarly, metformin suppressed Ang II – induced ROS accumulation in controls, whereas *Ppargc1a* deficiency not only abolished this antioxidant effect but also potentiated oxidative stress (Figure 7J-7K). Together, VSMC-specific PGC-1α loss disrupts mitochondrial integrity and bioenergetics, rendering cells refractory to metformin’s protective actions.

### 3.8 Metformin activates the AMPK – SIRT1 – PGC-1α axis to suppress vascular senescence; Dual Inhibition of AMPK and SIRT1 abrogates its protective Effects

Metformin activates AMPK through dose-dependent mechanisms: at higher concentrations, it inhibits mitochondrial complex I, elevating the AMP:ATP ratio and promoting AMPK phosphorylation ^43^. At lower (clinically relevant) doses, it engages a lysosomal AXIN/LKB1-dependent pathway independently of AMP/ATP fluctuations ^44^. Moreover, SIRT1 and AMPK frequently cooperate to suppress cellular senescence ^45,46^, and—in specific cellular contexts—AMPK activation can potentiate SIRT1 deacetylase activity via NAD^+^ elevation ^47^.

In AAA tissues, AMPK phosphorylation (Thr172), total AMPK, and SIRT1 were markedly reduced in the Model group, whereas metformin restored their expression (Figure 8A-D). To interrogate this axis, we administered metformin with either Compound C (CC; AMPK inhibitor) or Ex-527 (SIRT1 inhibitor) (Figure 8L). The aortic diameter was significantly enlarged in inhibitor-treated mice, demonstrating that AMPK or SIRT1 blockade abrogates metformin’s protective effect (Figure 8M-O). Mechanistically, CC suppressed AMPK Thr172 phosphorylation and SIRT1 expression, whereas Ex-527 selectively reduced SIRT1 without affecting AMPK phosphorylation (Figure 8E-H). Both inhibitors abolished metformin-induced PGC-1α upregulation (Figure 8E, I, P, S), supporting a functional AMPK– SIRT1–PGC-1α signaling module. Notably, VSMC-specific Ppargc1a deletion reduced both SIRT1 protein levels and the fraction of SIRT1 ⁺ VSMCs (Figure S12A-S12B), suggesting bidirectional positive regulation between PGC-1α and SIRT1.

**Figure 8.**
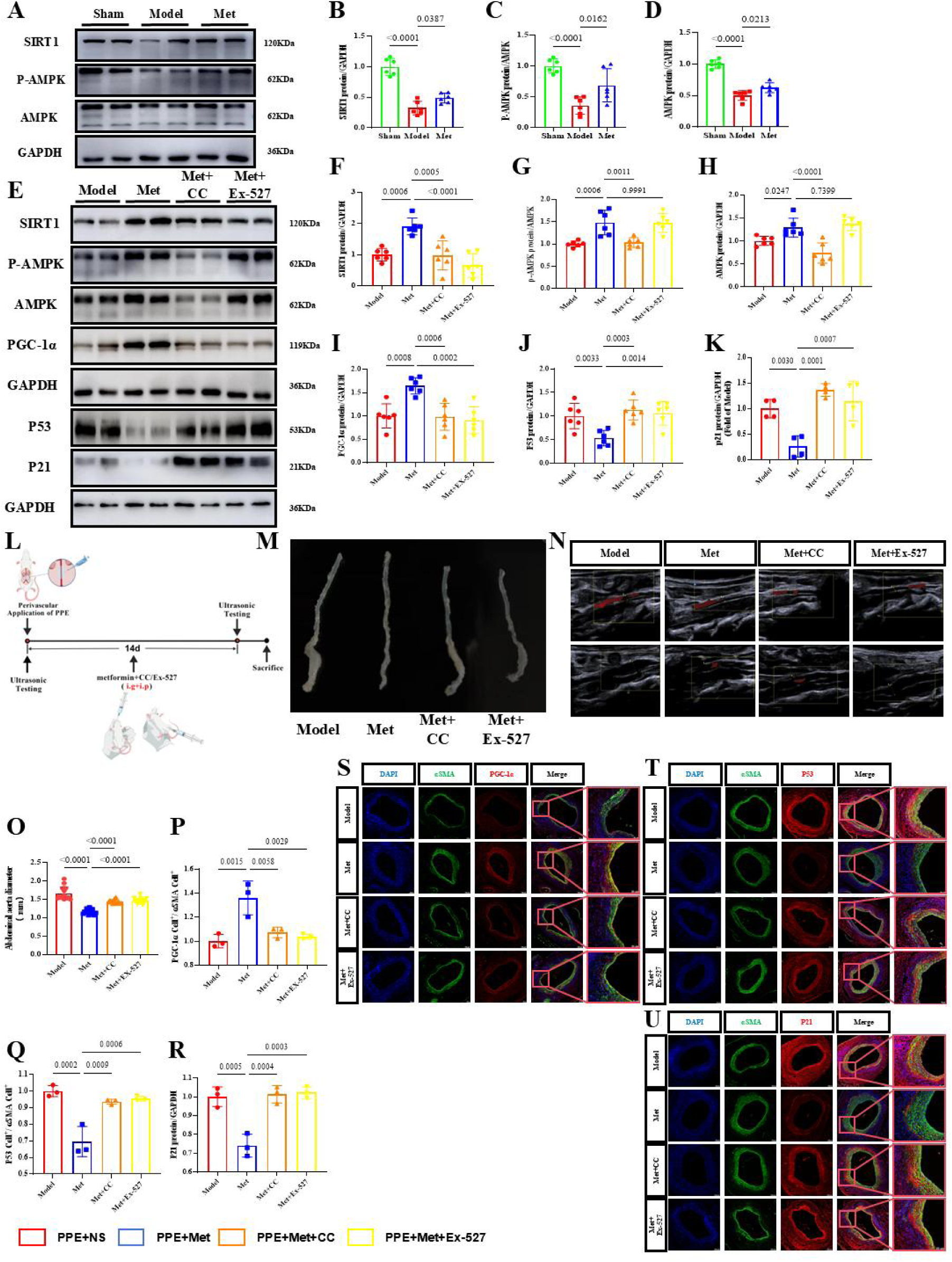
Metformin upregulates PGC-1α via the AMPK/SIRT1 pathway to counteract senescence. (A) Western blot of SIRT1, phosphorylated AMPK (p-AMPK), and total AMPK in aortic lysates. (B–D) Quantification of (A) (n = 6). (E) Western blot of SIRT1, p-AMPK, AMPK, PGC-1α, p53, and p21 in aortas from Model, Metformin (Met), Met+Compound C (CC, AMPK inhibitor), and Met+Ex-527 (SIRT1 inhibitor) groups. (F–K) Quantification of (E) bands (n = 6 for most; n = 4 for p53/p21 due to membrane stripping). (L) Experimental design: After AAA induction, mice received (i.g.) vehicle or metformin (50 mg/kg/day), combined with (i.p.) vehicle, CC (10 mg/kg/day), or Ex-527 (5 mg/kg/day). Vehicle formulation: 5% DMSO, 40% PEG300, 5% Tween-80, 50% saline. (M) Representative aortic morphology (left to right: Model, Met, Met+CC, Met+Ex-527). (N) Ultrasound images pre- and post-treatment. (O) Aortic diameter quantification from (N) (n = 16). (P–R) Quantification of PGC-1α-, p53-, and p21-positive cells from (S–U), respectively (n = 3). (S–U) Immunofluorescence for PGC-1α, p53, and p21. Data are mean ± SD. One-way ANOVA with Tukey’s post hoc test (B–D, F–K, O–R); *p* < 0.05 was considered significant.

Consistently, CC and Ex-527 reversed metformin’s suppression of senescence markers p53 and p21 (Figure 8E, J-K), and fully abolished its anti-senescence action in vivo, as evidenced by increased p53⁺/p21⁺ VSMCs in aortic tissuest (Figure 8Q-8R, 8T-8U).

### 3.9 Pharmacological Blockade of AMPK or SIRT1 Abrogates Metformin’s Protection Against VSMC Phenotypic Modulation in AAA

Given that either CC or Ex-527 abolished metformin’s anti-senescence action, we next examined their impact on VSMC phenotypic modulation. Consistent with this, CC or Ex-527 markedly elevated IL-6 and TNF-α in aortic tissues, reversing metformin’s suppression (Figure 9A-C). Immunofluorescence further showed increased IL-6 ⁺ and TNF-α^+^ cells in the media (Figure 9D-E, G-H), reflecting enhanced vascular inflammation. Critically, the rise in inflammatory cytokines coincided with a marked loss of the contractile marker SMTN (Figure 9F, 9I), supporting a link between vascular inflammation and VSMC dedifferentiation. Given the reduction in contractile markers (Fig. 9F, I), this inflammatory milieu likely contributes to, or results from, VSMC phenotypic modulation. Histologically, H&E staining demonstrated exacerbated immune cell infiltration upon inhibitor co-treatment (Figure 9J). Similarly, EVG and VB staining revealed severe elastic fiber fragmentation, in stark contrast to the protective effect of metformin monotherapy (Figure 9J).

**Figure 9.**
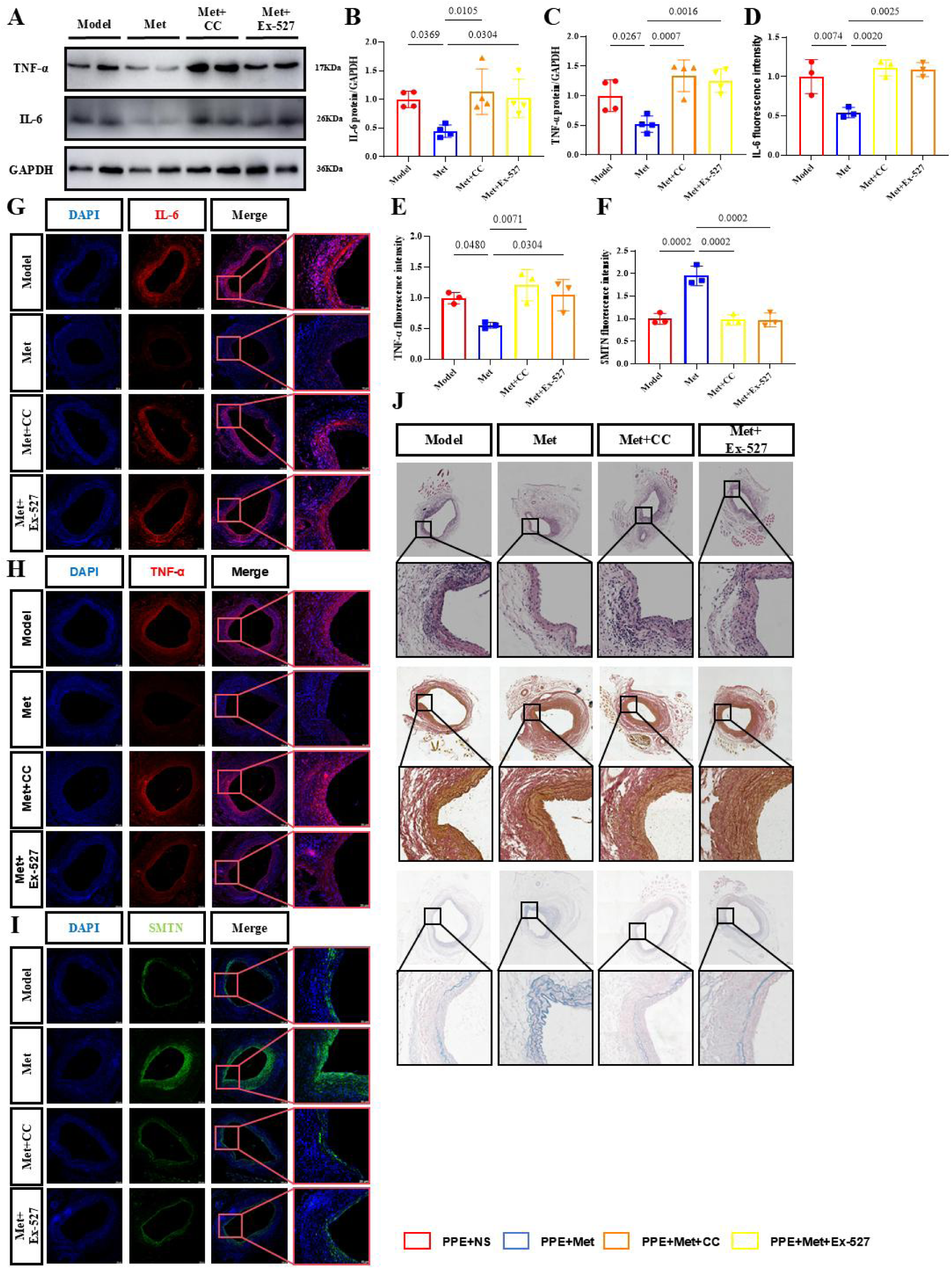
Dual inhibition of AMPK and SIRT1 abolishes metformin-mediated suppression of VSMC phenotypic switching. (A) Western blot of TNF-α and IL-6. (B, C) Quantification of (A) (n = 4). (D–F) Fluorescence intensity quantification for TNF-α, IL-6, and SMTN from (G–I) (n = 3). (G–I) Immunofluorescence staining. (J) H&E, EVG, and VB staining of aortic tissues (n = 3). Data are mean ± SD. One-way ANOVA with Tukey’s post hoc test (B, C, D, E, F); *p* < 0.05 was considered significant.

Collectively, the loss of metformin’s protective effect upon AMPK or SIRT1 inhibition—coupled with reduced PGC-1α (Fig. 8), SMTN, and enhanced inflammation—supports a model wherein the AMPK– SIRT1 – PGC-1α axis is critical for restraining pathological VSMC dedifferentiation in AAA.

## 4. Discussion

AAA is a life-threatening degenerative vascular disease for which no effective pharmacotherapies exits, with current management limited to surveillance and surgical repair. Epidemiological stdudies consistently associate metformin use with attenuated AAA expansion and reduced rupture risk ^12,13,48,49^, yet the underlying mechanisms remained elusive. Here, we provide mechanistic evidence that metformin interrupts a feed-forward loop linking VSMC senescence, phenotypic dedifferentiation, and vascular inflammation by restoring mitochondrial homeostasis via the AMPK – SIRT1 – PGC-1α axis.

Alghough VSMC senescence has been implicated in AAA pathogenesis ^50–52^, its functional coupling to mitochondrial dysfunction and phenotypic plasticity was unclear. Our genetic and pharmacological data establish VSMC-intrinsic PGC-1α as a pivotal and non-redundant regulator of vascular resilience: its specific deletion accelerates aneurysm progression and renders vessels refractory to metformin’s protection, highlighting mitochondrial integrity as a cell-autonomous determinant of VSMC fate.

Although PGC-1α’s role in mitochondrial maintenance is conserved across tissues ^53–56^, its specific function in vascular homeostasis has been underexplored. Our work identifies VSMC-intrinsic PGC-1α as a guardian against stress-induced dedifferentiation — its deficiency triggers mitochondrial fragmentation, bioenergetic collapse, and ROS overproduction, culminating in accelerated senescence. Critically, PGC-1α and SIRT1 exhibit mutual functional dependence in VSMCs: *Ppargc1a* deletion reduced SIRT1 expression (Fig. S11), whereas SIRT1 inhibition abrogated metformin-induced PGC-1α upregulation (Fig. 8). This interdependence forms a self-reinforcing module; consistent with this, its genetic or pharmacological disruption accelerates AAA progression and exacerbates key pathological features — including mitochondrial fragmentation, VSMC dedifferentiation, and medial inflammation—thereby shaping a distinct disease trajectory.

Mechanistically, metformin acivates AMPK primarily by inhibiting mitochondrial complex I, leading to increased AMP:ATP ratios and AMPK phosphorylation ^57,58^, This, in turn, elevates NAD ⁺ bioavailability, potentiating SIRT1-mediated deacetylation and activation of PGC-1α^47^ — a modification essential for its transcriptional co-activation. Notably, VSMC-specific *Ppargc1a* deletion reduced SIRT1 protein levels (Fig. S11), suggesting that — in this cellular context — PGC-1α may also support *Sirt1* expression, forming a reinforcing loop. Disruption of any node (via genetic KO or pharmacological inhibition) collapses the entire module, leading to mitochondrial failure and senescence. We thus propose a PGC-1α-centered feed-forward loop in AAA pathogenesis: initial stressors ↓ PGC-1α → ↓ SIRT1 activity → hyperacetylated/inactive PGC-1α → worsened mitochondrial dysfunction → amplified ROS/DNA damage → accelerated senescence. Metformin intervenes at the apex by boosting PGC-1α expression/activity, thereby resetting mitochondrial homeostasis.

Recent spatial and single-cell transcriptomic analyses have identified a distinct “inflammatory” VSMC subpopulation in human and murine AAA, characterized by elevated expression of chemokines (Ccl2, Ccl19) and adhesion molecules (Icam1, Vcam1) ^59,60^. This subset promotes immune cell recruitment and medial inflammation, and our data position PGC-1α deficiency as a key driver of this pathological transition.

In summary, our work establishes the AMPK– SIRT1 – PGC-1α– mitochondria axis as a central regulator of VSMC fate in AAA, through which metformin suppresses senescence and phenotypic dedifferentiation to restore vascular homeostasis, thereby providing a mechanistic rationale for its repurposing. While murine models cannot fully recapitulate human AAA heterogeneity, the conservation of this pathway supports its translational potential. Future studies should evaluate PGC-1α-targeted strategies— including SIRT1 activators (e.g., SRT2104), NAD ⁺ boosters, or mitochondria-directed antioxidants — in large-animal models and AAA patient-derived cells, paving the way for precision interventions that stabilize the vulnerable aortic wall.

### Affiliations

School of Rehabilitation Medicine, Gannan Medical University, Ganzhou, China (B. G.), School of Basic Medicine, Gannan Medical University, Ganzhou, China (Y. Z., Y. L., X. Y., D. Y., S. L., J. Z., B. L., Y. J., S. D., Z. Y.), Key Laboratory of Prevention and Treatment of Cardiovascular and Cerebrovascular Diseases, Ministry of Education, Gannan Medical University, Ganzhou, China (J. X, Z. Y.), Department of Cardiology, First Affiliated Hospital of Gannan Medical University, Ganzhou, China (J. X).

### Sources of Funding

This work was supported by the National Natural Science Foundation of China (grant no.82260288 and 82260104)

## Acknowledgments

The authors thank Dr. Lv Qing for his generosity in sharing the *Itga8*-*CreER*^T2^ mice.

