## Supplementary Figures for "Metformin Stabilizes the Abdominal Aorta in Aneurysm by Restoring VSMC Mitochondrial Homeostasis via the AMPK–SIRT1–PGC-1α Axis"

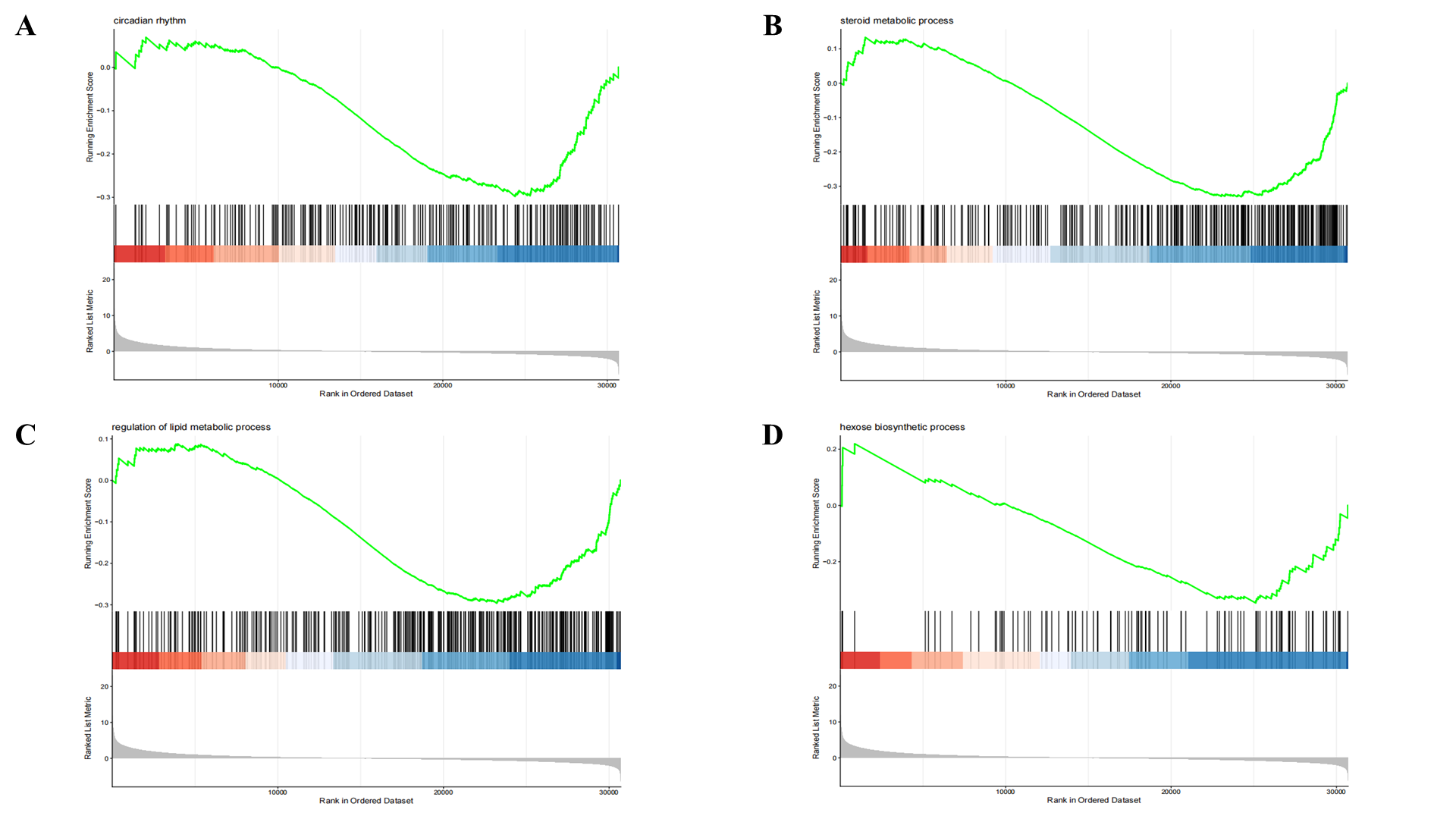
**Figure S1. GSEA of transcriptomic profiles in Sham vs. Model aortas (GEO database, hallmark gene sets).**
(A–D) Enrichment plots showing upregulated pathways in Model group.

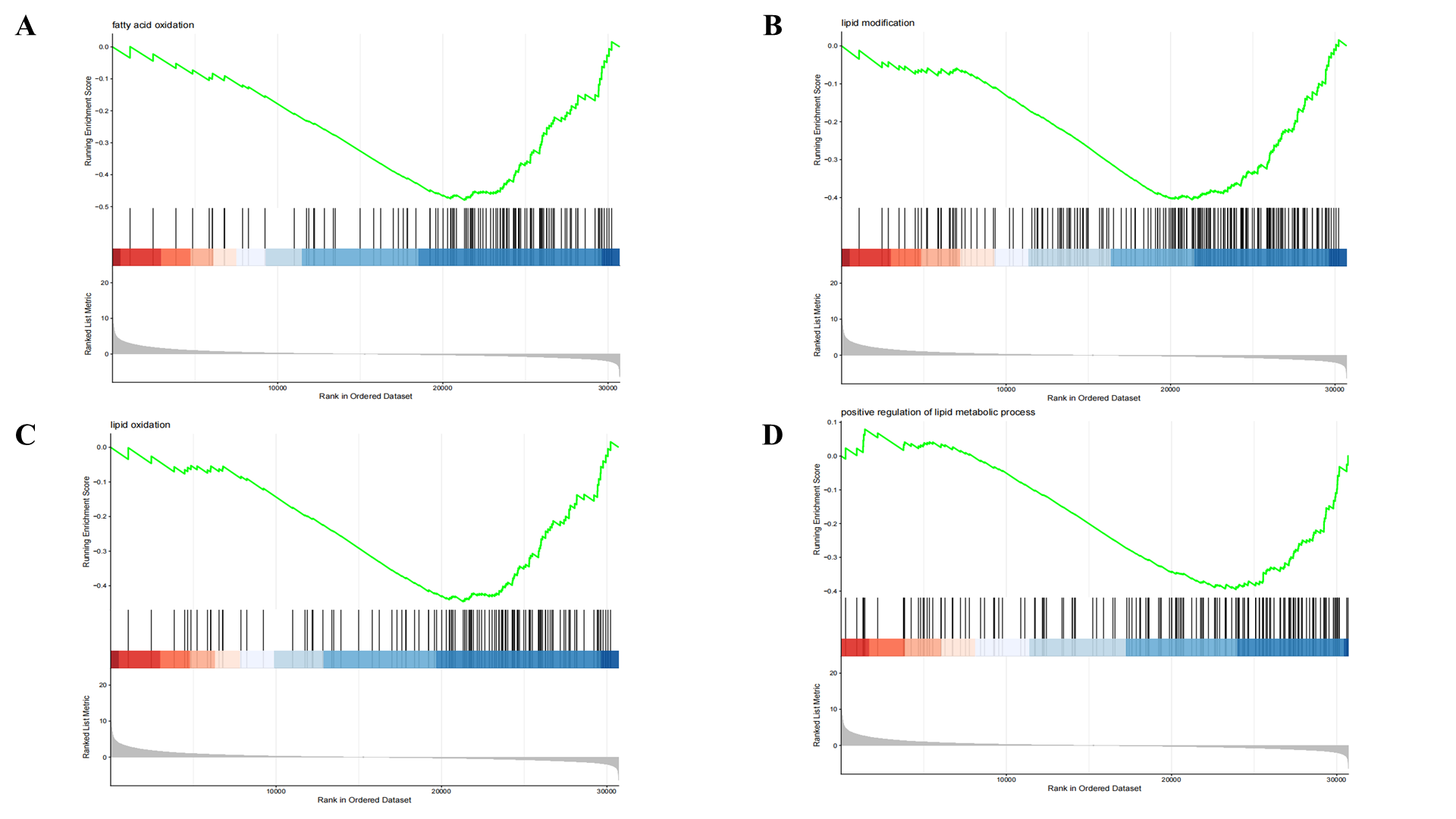

**Figure S2. GSEA of transcriptomic profiles in Sham vs. Model aortas (GEO database, C2 curated gene sets).**
(A–D) Enrichment plots indicating dysregulated biological processes.

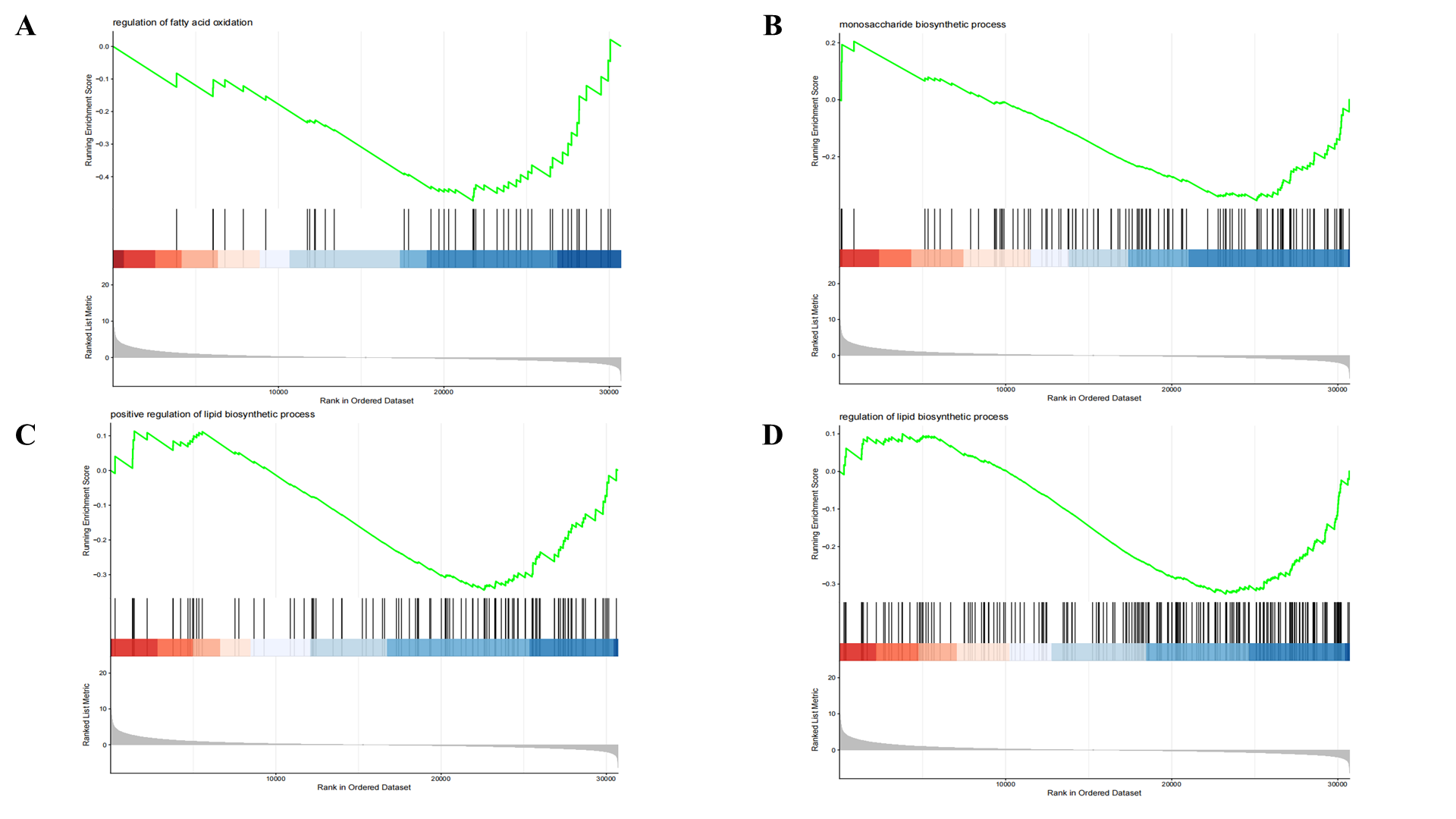

**Figure S3. GSEA of transcriptomic profiles in Sham vs. Model aortas (GEO database, C2 curated gene sets).**
(A–D) Enrichment plots indicating dysregulated biological processes.

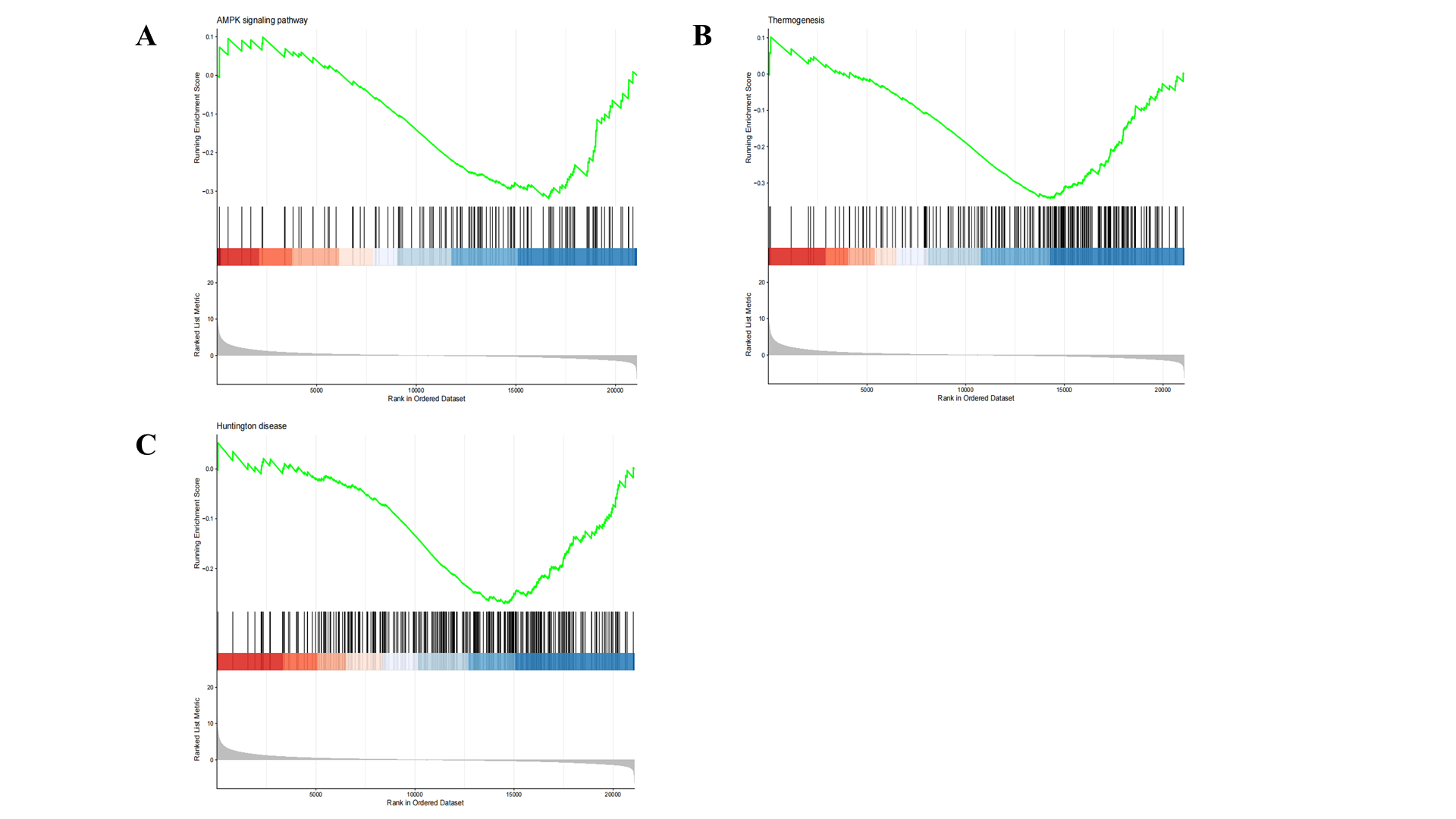

**Figure S4. GSEA based on KEGG pathways.**
(A–C) Key enriched KEGG pathways in Model vs. Sham aortas.

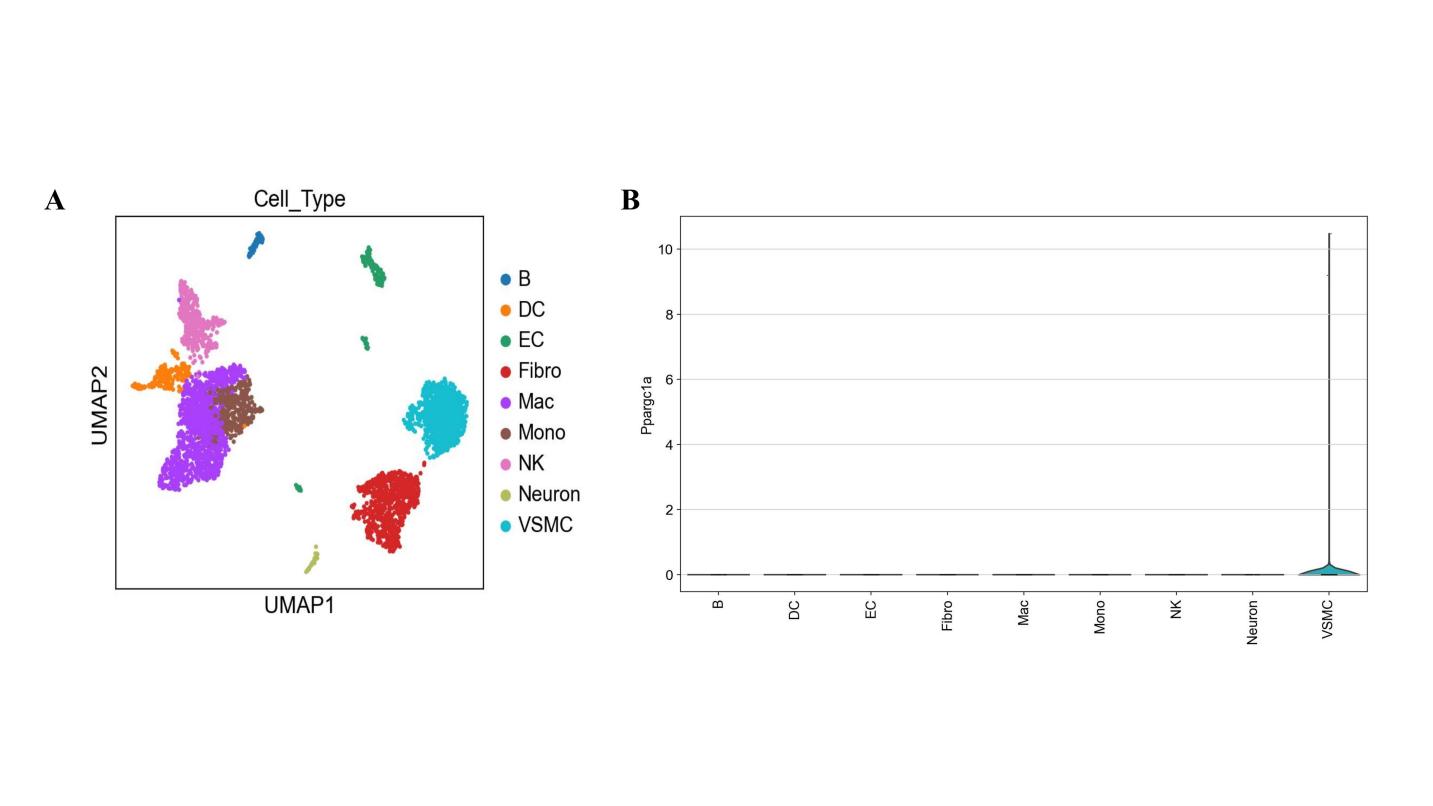

**Figure S5. Single-cell RNA sequencing of mouse abdominal aortas.**
(A) UMAP clustering of major aortic cell populations.
(B) *Ppargc1a* expression across annotated cell types.

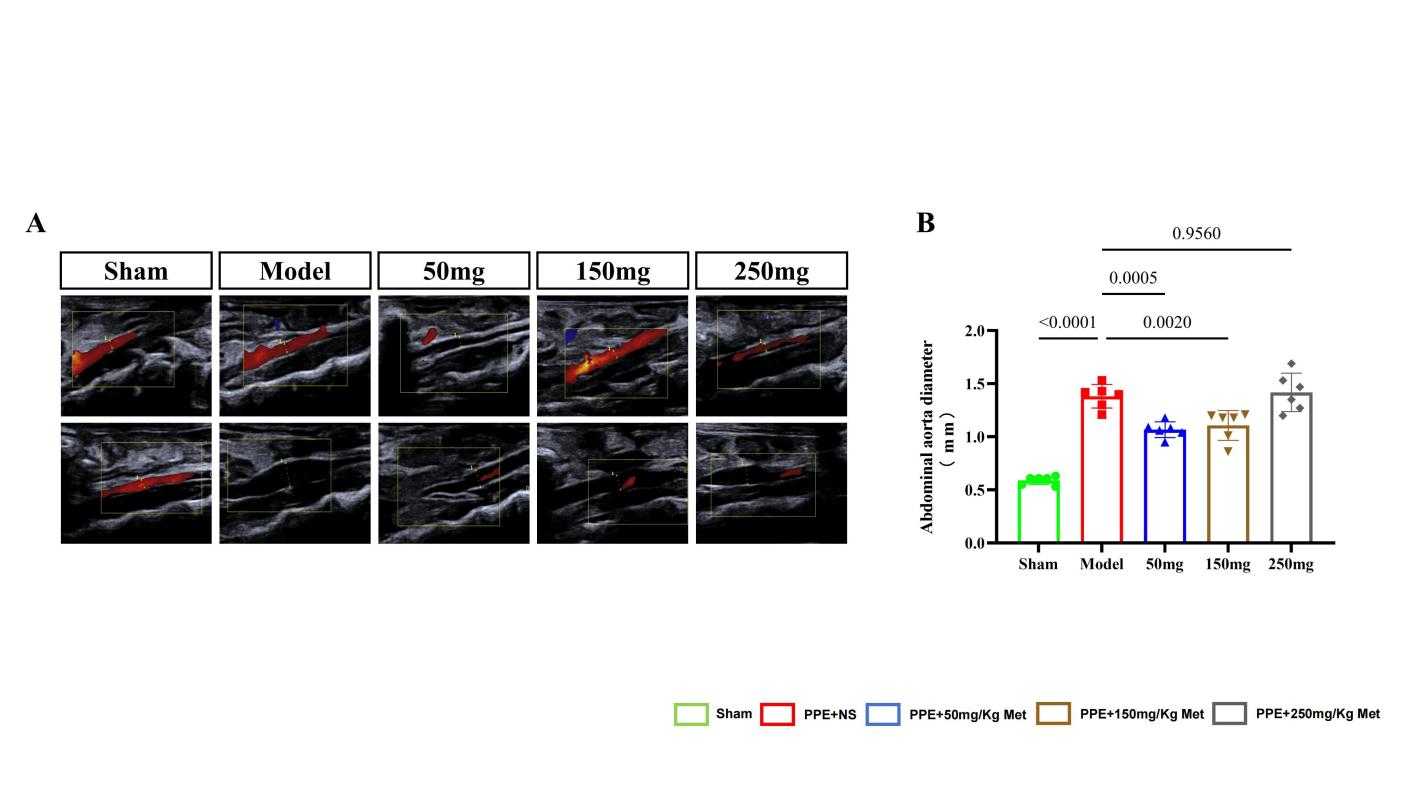

**Figure S6. Dose-response screening of metformin *in vivo*:**
(A) Serial ultrasound images.
(B) Maximal aortic diameter (n = 6/group). One-way ANOVA; *p* < 0.05.

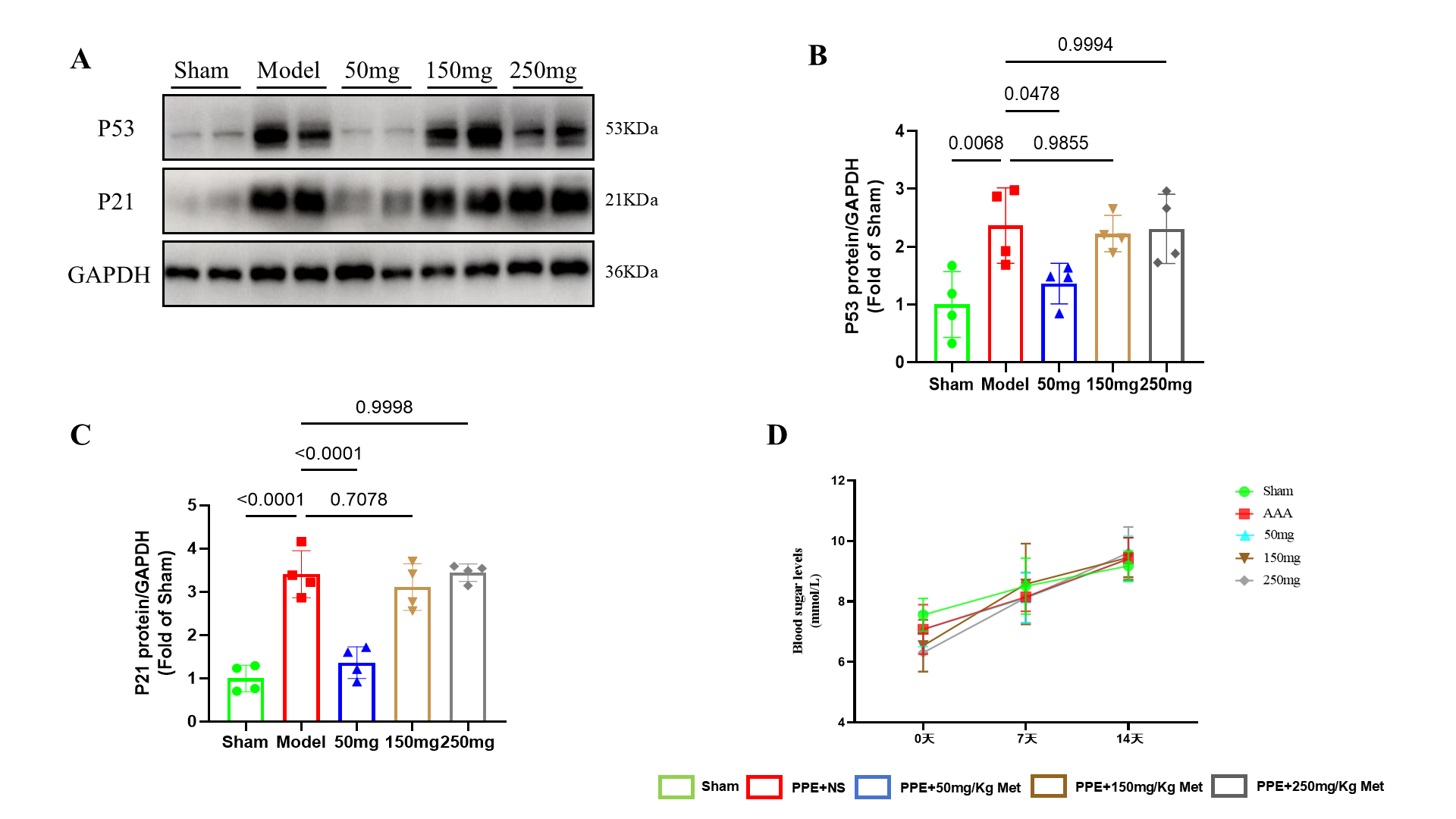
**Figure S7. Pharmacological dose optimization.**
(A) Western blot of p53/p21 in aortas from mice treated with 0, 50, 150, or 250 mg/kg metformin.
(B, C) Densitometry (n = 4).
(D) Fasting blood glucose at days 0, 7, and 14 (n = 6). One-way ANOVA (B, C); two-way repeated-measures ANOVA (D); *p* < 0.05.

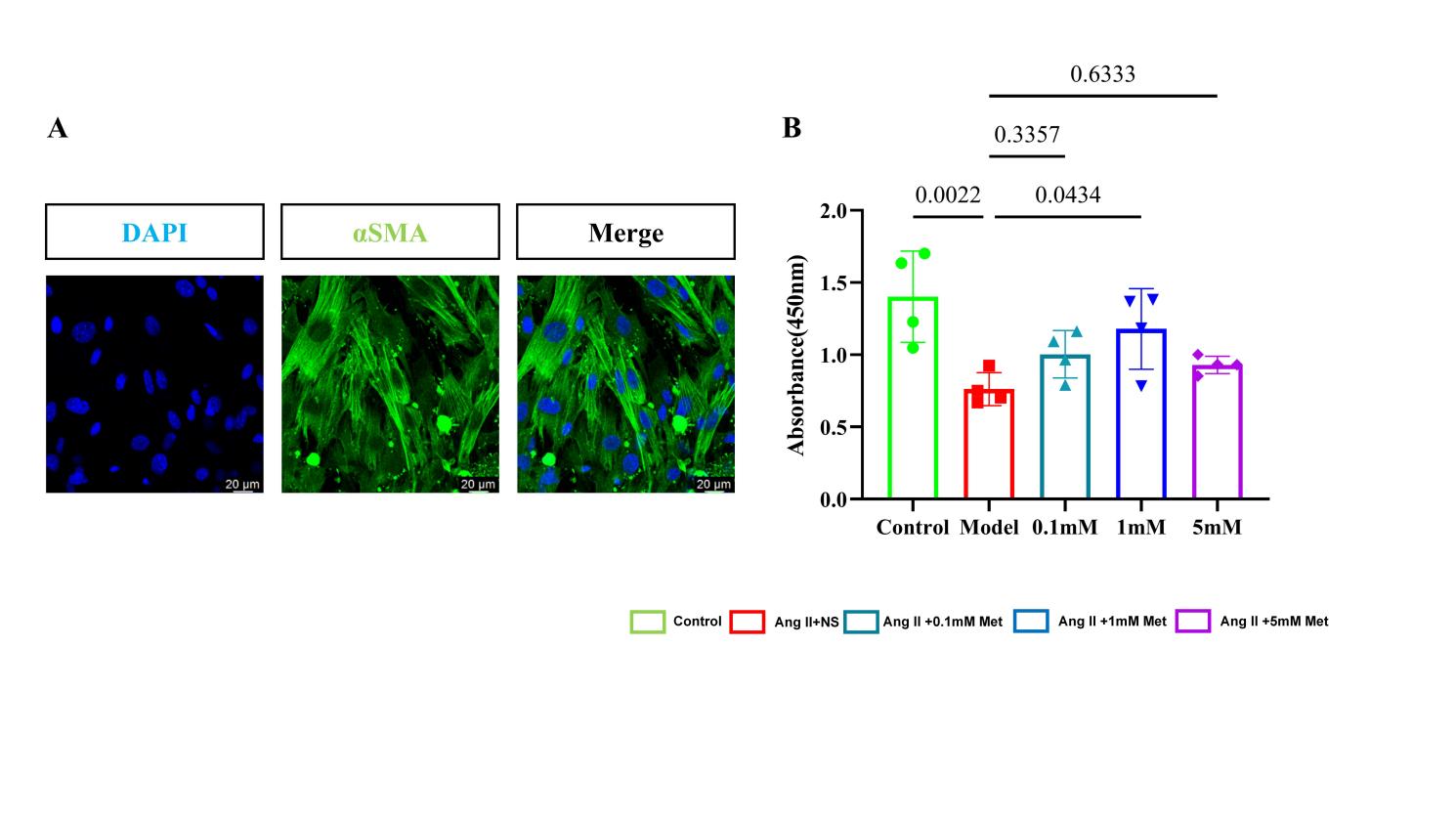

**Figure S8. *In vitro* drug screening in primary VSMCs.**
(A) α-SMA immunofluorescence confirming VSMC identity.
(B) CCK-8 assay of cell viability after exposure to 0.1, 1, or 5 mM metformin (n = 4). One-way ANOVA; *p* < 0.05.

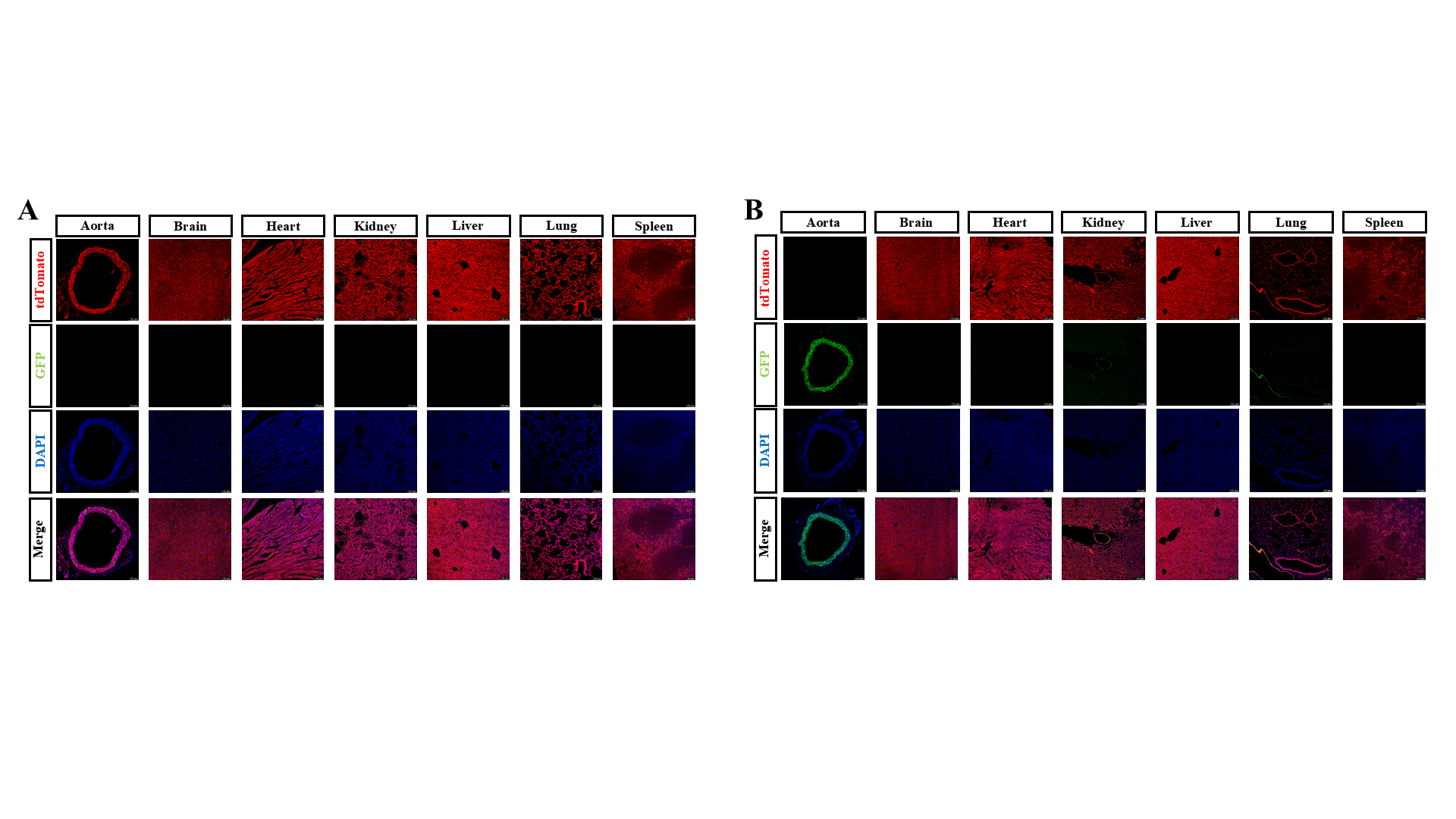

**Figure S9. Tamoxifen-induced recombination efficiency in *Rosa26*mTmG/+; *Itga8-Cre*+/− mice.**
(A, B) GFP (recombined) vs. tdTomato (non-recombined) fluorescence in aorta, brain, heart, kidney, liver, lung, and spleen.

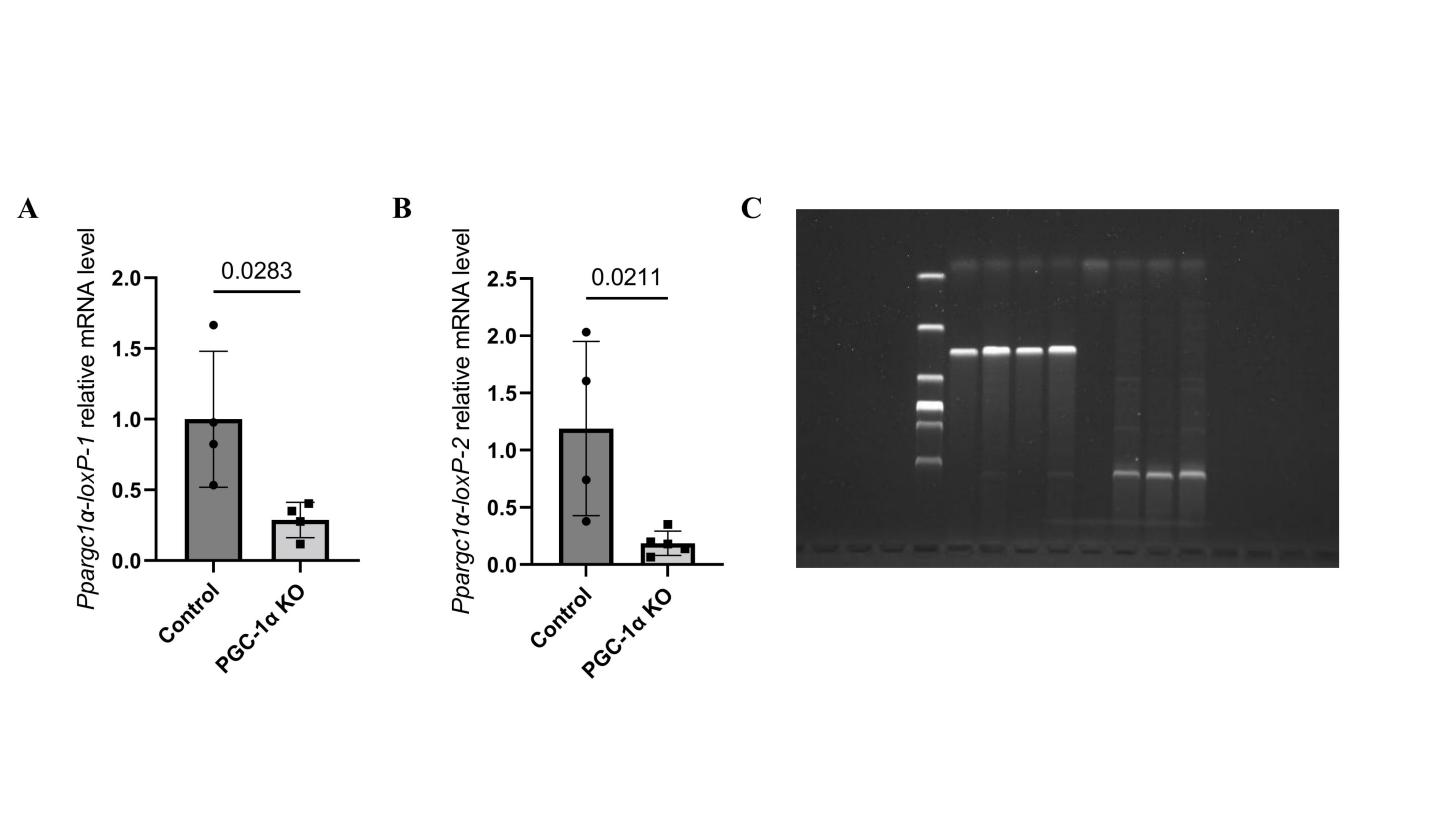

**Figure S10. Validation of *Ppargc1a* knockout in VSMCs.**
(A, B) qRT-PCR of *Ppargc1a* mRNA in aortic tissues (n = 4–5).
(C) Genomic PCR confirming excision of *Ppargc1a* floxed allele (left: KO, right: Ctrl). Two-tailed *t*-test (A, B); *p* < 0.05.

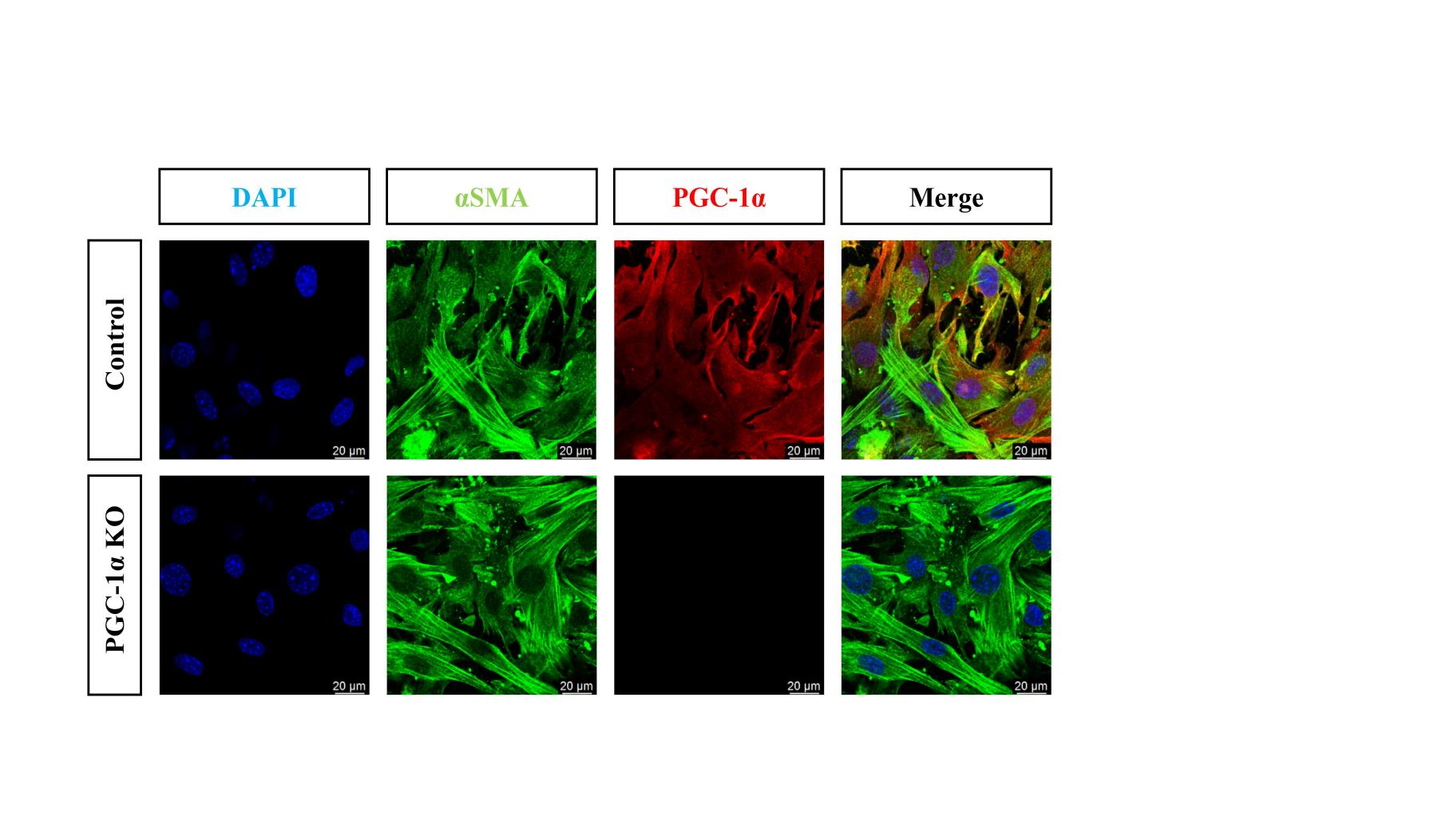

**Figure S11. Characterization of primary VSMCs and *Ppargc1a* KO efficiency.** Immunofluorescence co-staining of α-SMA (red) and PGC-1α (green); nuclei in DAPI.

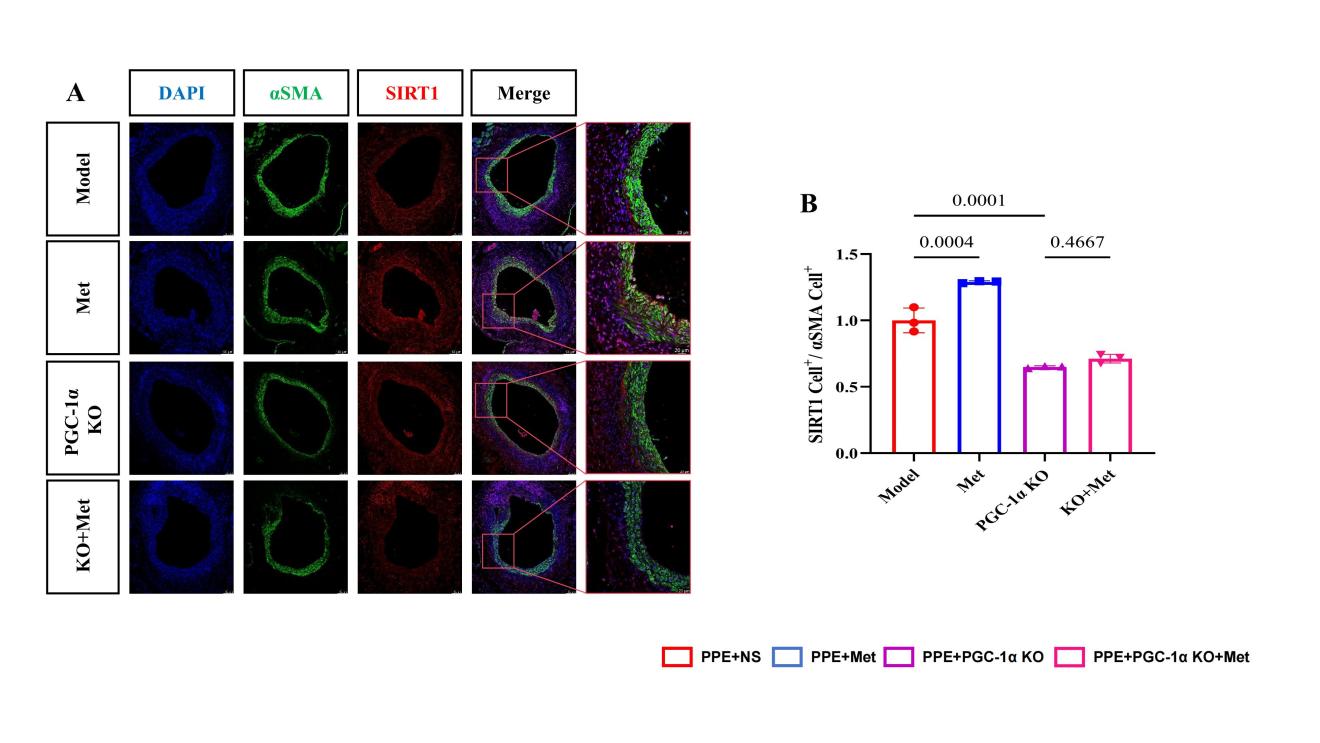

**Figure S12. SIRT1 expression upon *Ppargc1a* knockout.**
(A) Immunofluorescence of SIRT1 in aortic sections.
(B) Quantification of SIRT1-positive cells (n = 3). One-way ANOVA; *p* < 0.05.
